# The demographic history, genomic variation, and transcontinental genotype-phenotype-environment map of mungbean

**DOI:** 10.1101/2025.10.16.682767

**Authors:** Ya-Ping Lin, Hung-Wei Chen, Chih-Cheng Chien, Yu-Han Lien, Pei-Wen Ong, Chia-Min Yeh, Pei-Min Yeh, Min-Wei Chai, Colin A Douglas, AKM Mahbubul Alam, Mar Mar Win, Aditya Pratap, Venkata Naresh Boddepalli, Rael Karimi, Herbaud Phanuel Fortunat Zohoungbogbo, Jean Baptiste De La Salle Tignegre, James Y. Asibuo, Papias Hongera Binagwa, Emmanuel K. Mbeyagala, Shahid Riaz Malik, Maria G. Samsonova, Ji Ho Chu, Jo-Yi Yen, Eric Bishop von Wettberg, Ken Naito, Roland Schafleitner, Ramakrishnan M. Nair, Cheng-Ruei Lee

**Affiliations:** World Vegetable Center Headquarters, Tainan, Taiwan; Institute of Ecology and Evolutionary Biology, National Taiwan University, Taipei, Taiwan; Institute of Plant Biology, National Taiwan University, Taipei, Taiwan; Department of Agriculture and Fisheries, Queensland, Australia; Bangladesh Agricultural Research Institute (BARI), Gazipur, Bangladesh; Department of Agricultural Research (DAR), Nay Pyi Taw, Myanmar; ICAR-Indian Institute of Pulses Research (IIPR), Kanpur, India; World Vegetable Center South Asia /Central Asia, Telangana, India; Kenya Agricultural and Livestock Research Organization (KALRO, Katumani, Kenya; World Vegetable Center, West and Central Africa-Coastal and Humid Regions, Abomey-Calavi, Benin; World Vegetable Center, West and Central Africa-Dry Region, Bamako, Mali; CSIR-Crops Research Institute, Kumasi, Ghana; Tanzania Agricultural Research Institute (TARI) - Selian Centre, Arusha, Tanzania; National Agricultural Research Organization-National Semi-Arid Resources Research Institute (NARO-NaSARRI), Soroti, Uganda; Faculty of Agriculture and Animal Sciences (FAAS), Busitema University, Arapai Campus, Soroti, Uganda; National Agricultural Research Center (NARC) Islamabad, Pakistan; Peter the Great St.Petersburg Polytechnic University, Russia; National Agrobiodiversity Center, Jeonju-si, Jeollabuk-do, Republic of Korea; Vermont Agricultural Experiment Station, University of Vermont, Burlington, Vermont, USA; Research Center of Genetic Resources, National Agriculture and Food Research Organization, Tsukuba, Ibaraki, Japan; World Vegetable Center Mexico Office, CIMMYT Global Headquarters, Carretera México-Veracruz, Km 45, El Batán, 56237 Texcoco, México

**Keywords:** Mungbean, Domestication, Location Adaptation, Seed Coat Color, Genotype-Environment interaction

## Abstract

The breeding of mungbean, a crucial Asian legume, has been hampered by the lack of genomic resources. The International Mungbean Improvement Network (IMIN) aims to ensure global access to diverse germplasms and genomic resources. Using 780 worldwide wild and cultivated accessions, we report this species’ most comprehensive (pan)genomic variation, demographic history, and genotype-phenotype-environment map. Despite archaeological evidence of the earliest cultivation in South Asia, present-day wild populations only possess relict traces of shared polymorphisms with cultivars. We showed that parallel losses of black seed coats in two *Vigna* species were caused by the same mutational mechanism in the same gene. In large-scale cross-continent field trials, we found accessions from distant environments from the trial sites have lower performance, especially in high-heritability and high-yield sites, suggesting future breeding priority in benign conditions on accessions from similar environments. Our comprehensive genomic and trial resources facilitate future breeding success of this essential crop.

## Introduction

Mungbean (*Vigna radiata* (L.) Wilczek) is a short-duration legume (55–70 days) well-suited to tropical and subtropical regions, integrating efficiently into double- and inter-cropping systems, particularly with cereals. Mungbean enhances soil fertility and is a rich plant-based protein source with essential amino acids and micronutrients^1,2^. Mungbean benefits small-holder farmers due to its low input requirements and strong market demand, particularly in South and Southeast Asia^3^. However, limited genomic research, particularly in the correct linkage-map-based reference genomes and population genomics resources covering species-wide diversity, constrains genetic improvements^4–10^.

The International Mungbean Improvement Network (IMIN) was established in 2016 by the World Vegetable Center (WorldVeg) with support from the Australian Centre for International Agricultural Research. The network ensures global access to genetically diverse germplasm, including a mini-core collection of 289 accessions containing genetic variation of cultivars across environmental gradients of the world^11,12^. To date, IMIN has distributed the mini-core and conducted field trials across 23 gardens in 15 countries, resulting in 43 large-scale trials in 2016-2021 spanning 38°S to 36°N and covering extensive environmental gradients.

The first mungbean short-read genome (VC1973A) was available in 2014^13^ with later improvements^14^. A recent genome (JL7) was assembled with long-read technologies^15^. Breeders have observed that these assemblies do not fully correspond to the linkage maps^16^. While recent studies attempted genome-wide association studies (GWAS) and pan-genome investigation^15,17–20^, a comprehensive investigation of species-wide genomic variation, specifically incorporating wild and cultivated accessions across the species range, has yet to be conducted. The IMIN project, incorporating accessions encompassing comprehensive cultivar variation across the world^12^, is ideal for such a task.

Genomic studies investigating mungbean domestication were relatively few, likely due to the agronomic focus and lack of wild materials^21^. In a first attempt, we^22^ investigated the cultivar (*V. radiata* var. *radiata*, *radiat*a hereafter) and wild (*V. radiata* var. *sublobata*, *sublobata* hereafter) germplasms. For global cultivars, our recent study^12^ utilized >1000 accessions and uncovered a unique post-domestication expansion route, which was constrained by climatic conditions, resulting in cultivars with prolonged life history in the south of continental Asia to maximize yield and fast-cycling cultivars in the north for the short and dry growing season, at the cost of yield. Based on the assumption that crop accessions are locally adaptive, the “focused identification of germplasm strategy” identified germplasms originating from extreme environments as breeding parents^23^. However, for many crop accessions, little geo-reference information was available^24^. Since mungbean germplasms have strong gene-environment associations on the continental scale^12^, we aim to predict the local environments of >1,000 accessions and investigate the association with germplasm performance across the IMIN trial, shedding light on future breeding strategies.

In this study, the primary objectives were to: (1) assemble multiple high-quality reference genomes, (2) investigate species-wide genetic variation, (3) construct a comprehensive pan-genome, (4) identify loci associated with agronomic traits through large-scale IMIN trials, and (5) examine transcontinental genotype-by-environment (GxE) interactions. Combining genomics, population genetics, large-scale field trials, and the dissection of gene-environment associations, we aim to contribute to future breeding efforts of this critical crop.

## Results

### High-quality assemblies

We performed long-read sequencing using Oxford Nanopore Technologies (ONT) for one cultivated (Crystal) and two wild (CQ3649 and Karumbyar) accessions. Each accession yielded ∼70Gb of data, with an average read N50 exceeding 19 kb (Supplementary Table 1). Contigs were anchored onto 11 pseudo-chromosomes consistent with the linkage map (Supplementary Fig. 1, 2). Our assemblies exhibited fewer contigs with N50 of approximately 16 Mb (Table 1), surpassing previously published mungbean genomes (VC1973A and JL7)^14,15^.

**Table 1.** Statistics of mungbean genome assemblies.

| **Genomic feature** | **VC1973A<sup>14</sup>** | **JL7<sup>15</sup>** | **Crystal** | **CQ3649** | **Karumbyar** |
| --- | --- | --- | --- | --- | --- |
| **Species** | cultivar | cultivar | cultivar | wild | wild |
| **Total assembly size (Mb)** | 475.7 | 475.35 | 506.1 | 521.38 | 518.19 |
| **No. contigs** | 1511 | 632 | 203 | 200 | 277 |
| **Largest_contig (Mb)** | 12.73 | 30.2 | 38.94 | 36.68 | 39.34 |
| **Contig N<sub>50</sub> (Mb)** | 2.8 | 10.34 | 15.99 | 16.9 | 16.48 |
| **Scaffold N<sub>50</sub> (Mb)** | 47.1 | 43.79 | 45.7 | 46.74 | 45.05 |
| **percentage anchored to chr.** | 89.92 | 98.72 | 98.91 | 96.74 | 98.53 |
| **Number of gaps** | 1047 | 259 | 65 | 42 | 82 |
| **GC content (%)** | 33.27 | 33.45 | 33.36 | 33.6 | 33.1 |
| **Complete BUSCOs (%)** | 91.36 | 98.02 | 98.57 | 98.27 | 98.82 |
| **Protein coding genes** | 30,958 | 40,125 | 33,487 | 33,674 | 33,483 |

Comparing the synteny among adzuki bean (*Vigna angularis*), ours (cultivar: Crystal, wild: CQ3649 and Karumbyar), and previous assemblies (JL7 and VC1973A), high collinearity exists among adzuki and all genomes we assembled (Supplementary Fig. 3). We named and oriented our chromosomes following a recent study of multiple *Vigna* genomes^25^, facilitating future comparative genomics. Rearrangements existed between our assembly and JL7 in chromosomes 3, 6, 7, and 11, and ours was supported by the linkage maps (Supplementary Fig. 3). Repetitive sequence and gene density distribution indicated that Crystal correctly captured the chromosomal structure, with repeat-rich and gene-sparse regions located near the center of most chromosomes, suggesting centromeric regions (Supplementary Fig. 4).

### Population structure and demographic history

This study includes the most comprehensive collection capturing global mungbean genomic variation (Supplementary Table 2), including 168 wild (*sublobata* from Africa, South Asia, Continental and Island Southeast Asia, and Australia) and 612 cultivated accessions (*radiata*, advanced breeding lines and landrace diversity across traditional mungbean cultivation zones^12^). Four accessions of the closely related *Vigna mungo* were used as the outgroup. The sequencing depth exceeded 20x per accession, with an average mapping rate of 98.71% ± 2.40%. A total of 38,353,744 (38.4 million) bi-allelic SNPs were identified.

ADMIXTURE revealed low cross-validation error values at *K* = 2, 5, and 8 (Supplementary Fig. 5). At *K* = 2, accessions were separated into wild and cultivar groups (Supplementary Fig. 6). At *K* = 5, three wild groups exist: eastern Australian, western Australian (including Indonesia), and Asian-African accessions. Two groups exist for the cultivars (Supplementary Fig. 6). At *K* = 8, the wild accessions contain six geographically structured groups: South Asia (SubAS_SA), Southeast Asia (SubAS_SEA), Africa (SubAF), Indonesia (SubIDN), eastern Australia (SubAUe), and western Australia (SubAUw) (Fig. 1a). Two cultivar groups (Rad1 and Rad2) remained.

**Fig. 1.**
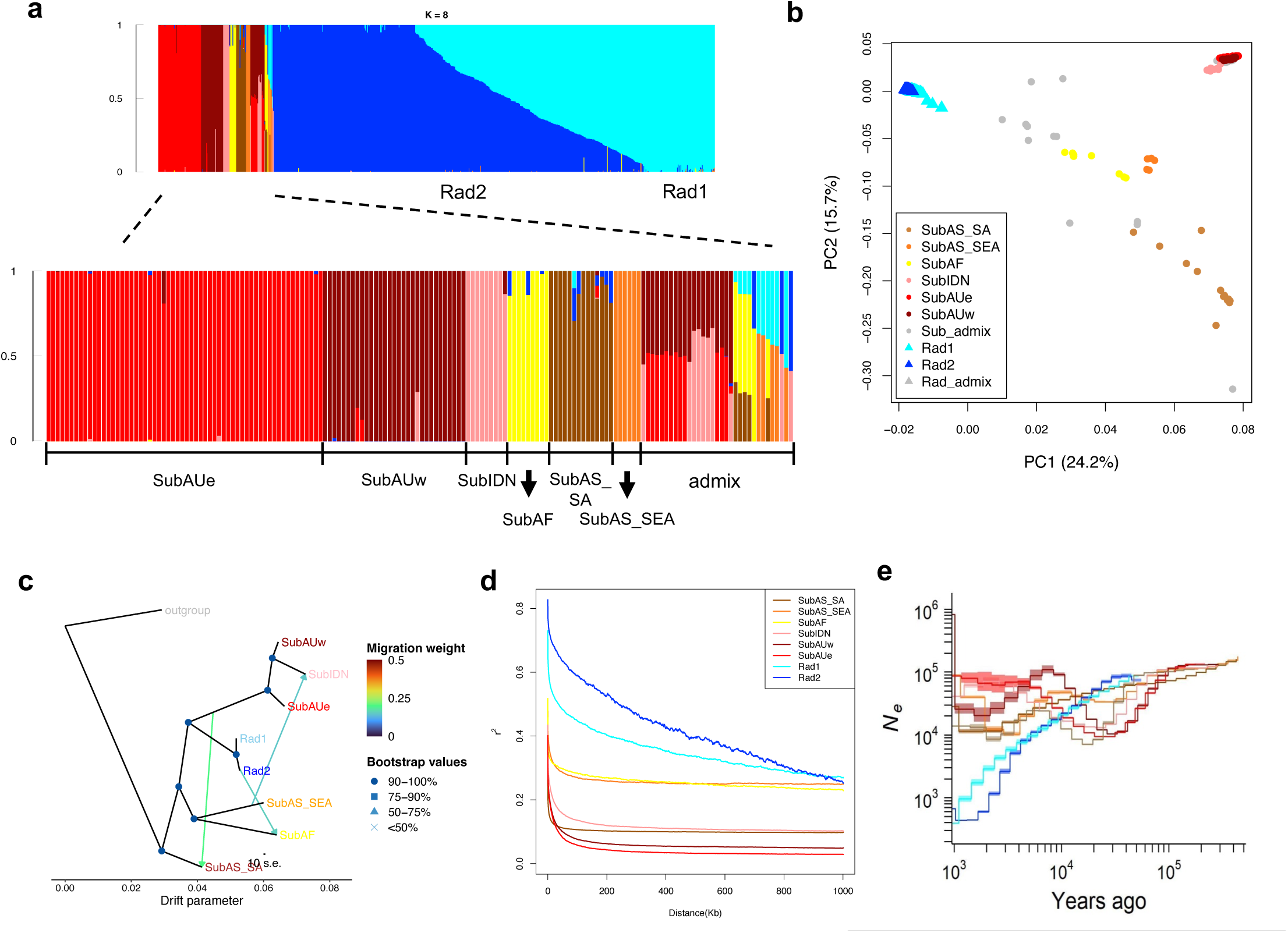
Species-wide genomic variation and demographic history of wild and cultivated mungbean. **a,** Population structure of K=8 estimated with ADMIXTURE. SubAS_SA: South Asian wild group (brown), SubAS_SEA: Southeast Asian wild group (orange), SubAF: African wild group (yellow), SubIDN: Indonesian wild group (pink), SubAUe: eastern Australian wild group (red), SubAUw: western Australian genetic groups (dark red), Rad1: cultivar group I (skyblue), Rad2: cultivar group II (blue). **b,** Principal component analysis (PCA) of the nuclear genome. **c,** Population-level phylogeny and direction of gene flow events among mungbean genetic groups using TreeMix. **d,** Decay of linkage disequilibrium. **e,** Population demography estimated with MSMC. The genetic group colors are consistent across all panels.

Most species closely related to mungbean existed in Asia, suggesting an Asian origin of mungbean^26^. Indeed, South Asian *sublobata* is the most diverged group from all others in principal component analysis (PCA, Fig. 1b) and TreeMix (Fig. 1c).

Southeast Asian and African groups shared a closer relationship (Fig. 1c and Supplementary Fig. 7). Another clade descended later, with the cultivars and Indonesian-Australian *sublobata* as sister clades. The individual-level maximum-likelihood tree is largely consistent (Supplementary Fig. 8). Among all groups, the highest nucleotide diversity existed in South Asian *sublobata* (Supplementary Fig. 9), and its linkage disequilibrium decays rapidly (Fig. 1d), supporting South Asia as the likely origin of wild mungbean.

Population demography estimated by MSMC showed a bottleneck for the wild groups likely coinciding with the previous glacial periods, while the cultivar groups experienced a continuous decline (Fig. 1e). Pairwise divergence times roughly reflected the order of divergence from TreeMix. In general, South Asian *sublobata* has the highest divergence time to all other clades (Supplementary Table 3 and Supplementary Fig. 10), and the three major *sublobata* clades (South Asia, Southeast Asia-Africa, and Australia) diverged at least 90 thousand years ago (kya, at relative cross-coalescent rate RCCR 0.75) to 70 kya (RCCR at 0.5). Notably, despite the distant relationship on TreeMix (Fig. 1c), *radiata* had the closest divergence time (< 10 kya) to the South Asian than to any other *sublobata* groups, likely reflecting a complex relationship that a bifurcating tree could not resolve.

### Domestication origin and gene flow

While archaeological evidence suggested that mungbean was domesticated in South Asia at about 4 kya^27^, TreeMix suggested South Asian *sublobata* was highly divergent to all groups. To validate TreeMix inferences, we calculated the pairwise *F_ST_*, *d_xy_*, and outgroup *f_3_* statistics (using *V. mungo* as the outgroup). While pairwise *F_ST_* might be affected by the within-population polymorphism, both *d_xy_* and outgroup *f_3_* showed that, among all wild groups, South Asian *sublobata* is the most distant to the cultivars, confirming TreeMix inferences (Supplementary Table 4,5).

Interestingly, the results showed that cultivars have higher genetic affinity (lowest *d_xy_* and highest outgroup *f_3_*) to the African and Indonesian *sublobata* (Supplementary Table 4,5). The genetic affinity is explained by the traces of gene flow from cultivars to African *sublobata* in TreeMix (Fig. 1c), and the ABBA-BABA tests based on the phylogeny (((SubAS_SEA, SubAF), RAD), *V. mungo*) obtained significantly positive values (*Z* = 6.0, *P* value < 1×10^−8^, Supplementary Table 6). This may be associated with the Austronesian migration bringing cultivated mungbean to East Africa^28^, which likely admixed with local *sublobata*. Consistently, from our recent study^12^, many present-day East-African cultivars have a Southeast Asian genetic background, supporting this hypothesis. While not detected by TreeMix, ABBA-BABA tests of (((SubAU, SubIDN), RAD), *V. mungo*) obtained significantly positive values (*Z* score > 3, *P* value < 1×10^−3^, Supplementary Table 6), supporting gene flow from cultivars into Indonesian *sublobata*. Despite the present large-scale mungbean cultivation in Australia, Australian *sublobata* did not possess stronger traces of cultivar gene flow than Indonesian *sublobata*, which was also observed from plant phenotype in the field^29^. Finally, another gene flow event was detected from continental Southeast Asian to Indonesian *sublobata* (Fig. 1c), consistent with their geographic proximity.

What might explain the archaeology-genetics inconsistency of mungbean domestication? During cultivation range expansion, suitable habitats might have been adopted to farmlands, and gene flow from the large cultivar population might dilute the wild gene pool^29^. In some crops, the closest wild progenitors were likely extinct, leaving all present-day wild groups highly distant from cultivars^30^. Given the lack of recombination in the chloroplast genome, one may still identify the wild haplotypes closest to mungbean cultivars. Individual-level neighbor-joining tree of the chloroplast shows a similar pattern to the nuclear genome (Supplementary Fig. 11): South Asian *sublobata* chloroplasts are the most divergent, followed by the splitting between other *sublobata* groups and the cultivars. Interestingly, several Indian *sublobata* accessions possessed chloroplast haplotypes closest to all cultivars, as evident from the haplotype network (Fig. 2a). This is less likely due to gene flow of cultivar haplotypes into wild accessions, since no existing cultivars possess the same haplotype as these Indian *sublobata*. Therefore, the relic chloroplast haplotypes from the closest wild progenitors likely still existed in India. Finally, despite archeological evidence suggesting two independent traces of early mungbean cultivation^27^, the chloroplast haplotype network and our recent study of a pod non-shattering gene^22^ suggest a single genetic origin of present-day mungbean cultivars.

**Fig. 2.**
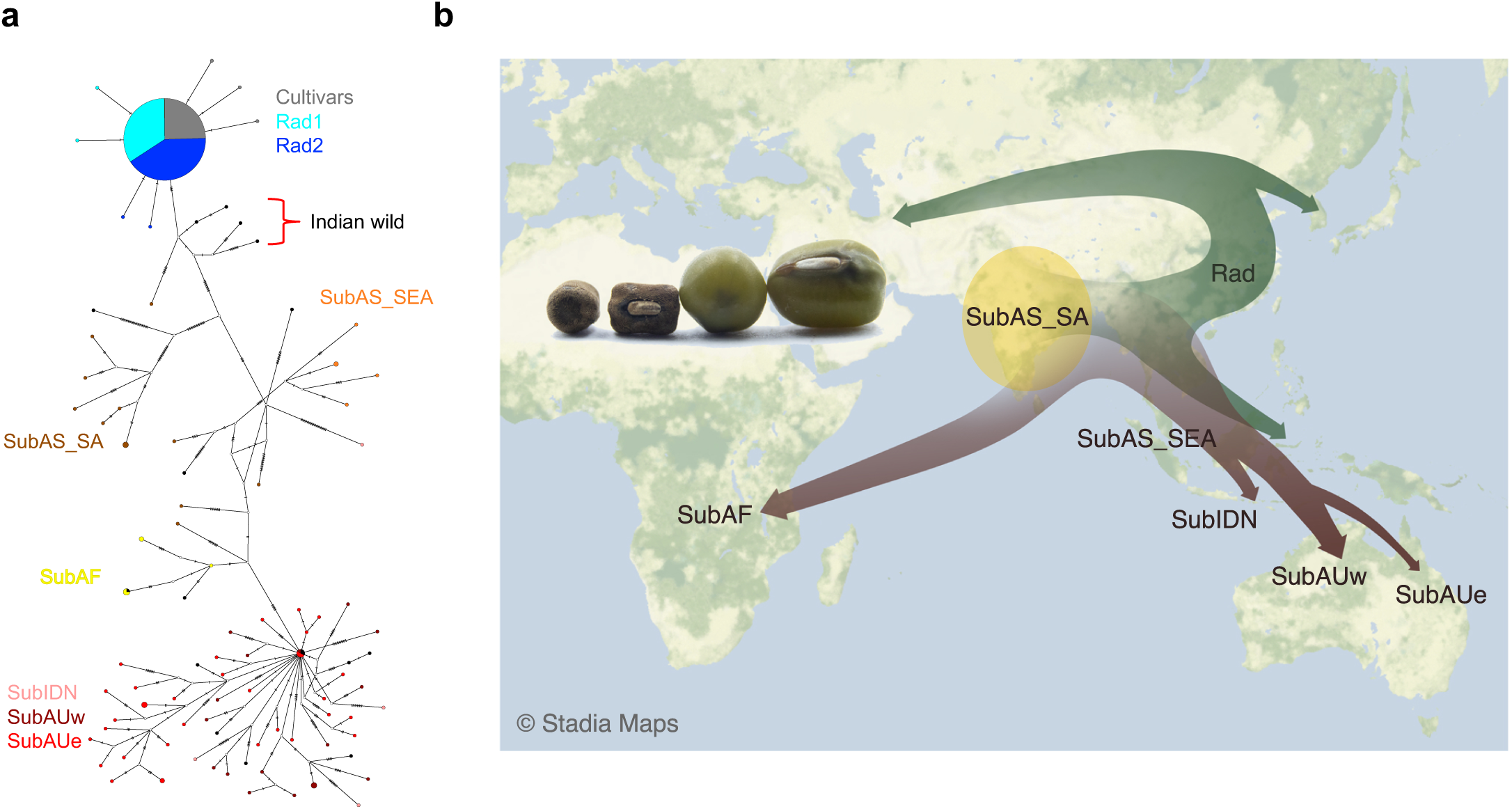
The origin and expansion of wild and cultivated mungbean. **a,** Chloroplast haplotype network. The nuclear-genome genetic group names and their colors were based on ADMIXTURE K=8 in Fig. 1a. The wild mungbean (although slightly admixed in their nuclear genomes) with the closest chloroplast haplotype to the cultivars were all from India, supporting the South-Asian domestication origin. **b,** Proposed route of mungbean range expansion based on the nuclear and chloroplast genome analyses.

Traces of South-Asian domestication might be inferred from patterns of shared SNPs. Excluding African and Indonesian *sublobata*, which had evident traces of gene flow with cultivars (Fig.1c & Supplementary Table 6), cultivars shared more unique SNPs (those only exist in cultivars and one specific *sublobata* group, Supplementary Fig. 12) with South Asian (91k, 1.34% among 6813k SNPs in South Asian wild) than with the others (Southeast Asia: 61k, 1.23% among 5025k; Australia: 65k, 1.25% among 5266k). ABBA-BABA tests revealed stronger gene flow of South Asian *sublobata* with other *sublobata* groups than *radiata* (Supplementary Table 6), suggesting the *radiata*-South Asian *sublobata* SNP sharing is less likely due to post-divergence gene flow. Our findings supported the hypothesis that wild mungbean originated in South Asia and subsequently spread to Africa, Southeast Asia, and Australia. Genomic traces of South-Asian domestication could be uncovered, and after domestication, mungbean cultivars followed a unique expansion route across Asia, as suggested by our previous studies^12^ (Fig. 2b).

To investigate the biological functions associated with selection during domestication and improvement, we performed GO enrichment analysis of candidate genes located in the top 5% intersection regions of XP-CLR and reduction of diversity tests. We identified 1,090 genes between wild and landrace accessions and 279 genes between landraces and improved lines (Supplementary Table 7 and 8). In the wild-landrace comparison, enriched GO terms were mainly related to terpenoid metabolism and protein/peptide transport (Supplementary Fig. 14a), suggesting the importance of secondary metabolite biosynthesis and intracellular trafficking during initial domestication. In contrast, the landrace-cultivar comparison revealed enrichment for macromolecule metabolism, protein modification/lysis, and RNA processing (Supplementary Fig. 14b), suggesting that improvements after domestication likely involved regulatory pathways and post-translational modification processes.

### The same mutation for the parallel loss of mottled black seed coats

During the domestication of most legumes, cultivars lost the mottled black spots on seed coats (Fig. 3a). In the adzuki bean (*Vigna angularis*), our study^31^ showed that this transition was caused by the deletion of an R2R3-MYB transcription factor gene, *VaPAP1*. The *PAP* locus consists of three homologs in wild adzuki, and the non-allelic homologous recombination between the flanking *VaPAP2a* and *VaPAP2b* deleted *VaPAP1*, leaving the chimeric *VaPAP2* gene in the cultivar genome^31^. Similar mottled black spot variation exists among mungbean accessions, and GWAS identified a peak corresponding to the *PAP* locus (Fig. 3b,c). Mungbean had a very similar mechanism: In both wild genomes, the *VrPAP1* gene was flanked by four *VrPAP2*s, and the non-allelic homologous recombination between the two outermost *VrPAP2* copies caused *VrPAP1* deletion in the cultivar (Fig. 3d).

**Fig. 3.**
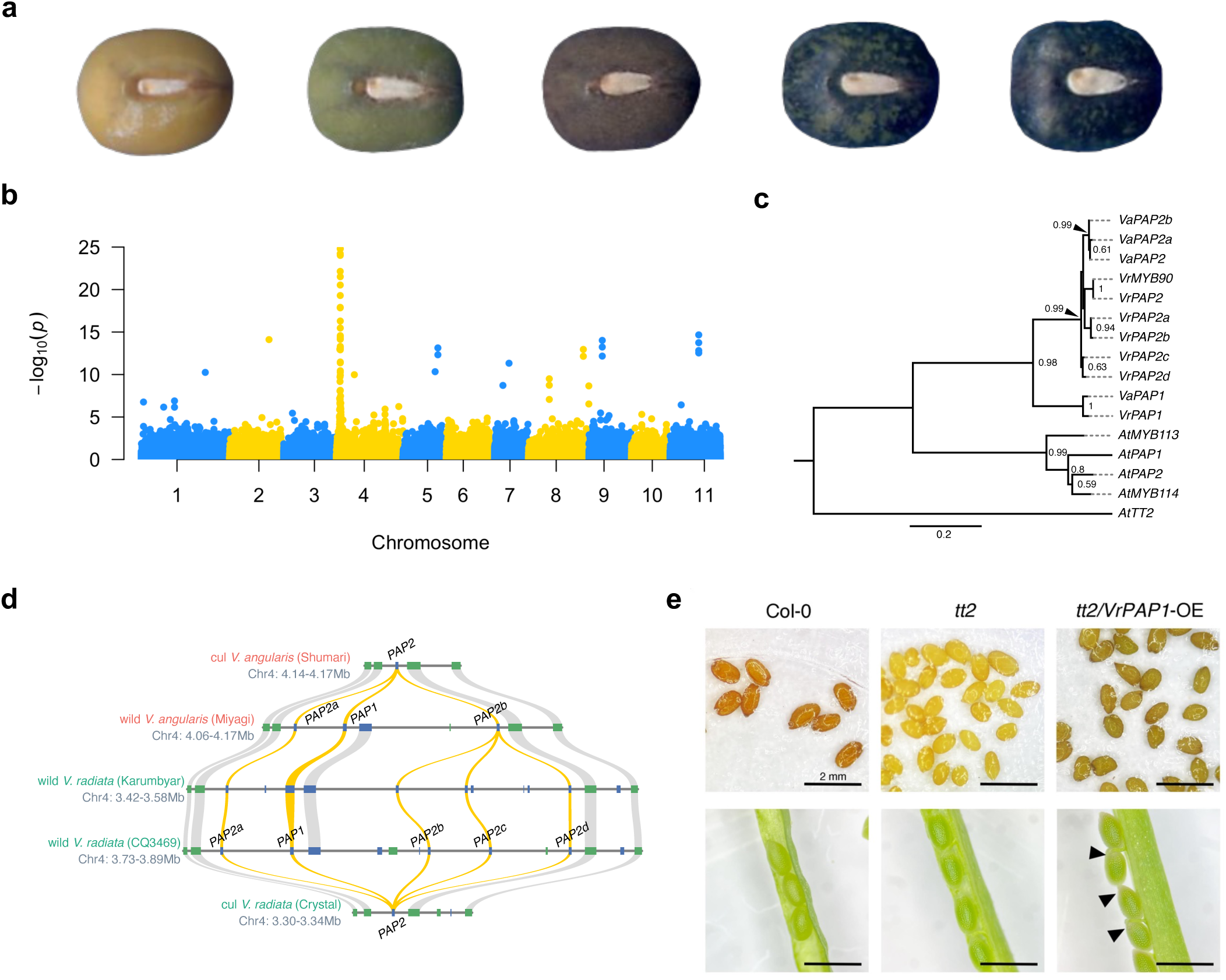
Parallel loss of dark seed coats during the domestication of adzuki and mungbean, caused by a similar type of mutation in the same gene. **a,** Seed coat color diversity of mungbean. **b,** Manhattan plot of genome-wide association study of mungbean seed coat color. The peak in chromosome 4 corresponds to the *PAP* locus, homologous to the locus resulting in the loss of dark seed coats in adzuki bean. **c,** Phylogeny of *PAP* homologs in mungbean (Vr: *Vigna radiata*), adzuki bean (Va, *Vigna angularis*), and *Arabidopsis thaliana* (At). **d,** Synteny plots of *PAP* gene clusters in three mungbean (below) and two adzuki bean (above) genomes. In both species, non-allelic homologous recombination between the two outermost *PAP2* homologs resulted in the deletion of *PAP1* in the cultivars, losing the dark pigmentation. **e,** Mature and young seeds of *Arabidopsis* Col-0 wildtype, *tt2* mutant line, and over-expression of *VrPAP1* in the *tt2* background.

*Vigna PAP* genes are closest to anthocyanin-associated *MYB*s in *Arabidopsis thaliana* (Fig. 3c), among which *AtMYB123* (*AtTT2*) was expressed in seed coats and resulted in lighter seed coats when mutated. When over-expressed in the *Attt2* mutant background, *VrPAP1* successfully restored the dark seed coat color (Fig. 3e), similar to *VaPAP1*^31^. Noticeably, while *VaPAP1* and *VrPAP1* were over-expressed across the whole plant body, anthocyanin accumulation was only observed in the seed coats but not in other plant parts. In contrast, Lin *et al*.^32^ overexpressed *VrMYB90* (the cultivar *VrPAP2* copy, Fig. 3c), whose expression was not detected in mungbean seeds, and found high anthocyanin accumulation across the whole plant. Therefore, the results suggested that *PAP1* and *PAP2* have diverged in their protein functions and tissues expressed, leaving *VrPAP1* as the candidate for the loss of mottled black seed coats during mungbean domestication. Our results demonstrated that the same genes and mutational mechanisms caused the parallel phenotypic evolution.

### Comprehensive species-wide pan-genome

Beyond the reference genome Crystal, the pan-genome comprises 100,533 contigs, with a total length of ∼267Mb and N50 of 4,584bp, containing 23,917 genes. Among the 780 accessions, we identified >57k genes, with 26,775 core and 30,292 dispensable genes. Compared to the pan-genome from 217 Chinese accessions^15^, with >43k genes sufficiently represented by <20 accessions, our comprehensive species-wide analyses identified more genes whose count did not saturate until >400 accessions were considered (Fig. 4a). The results are consistent with the current and previous studies that wild mungbeans possessed much higher genetic variation than cultivars^22^ and that East Asian cultivars possessed relatively low genetic variation among cultivar groups^12^.

**Fig. 4.**
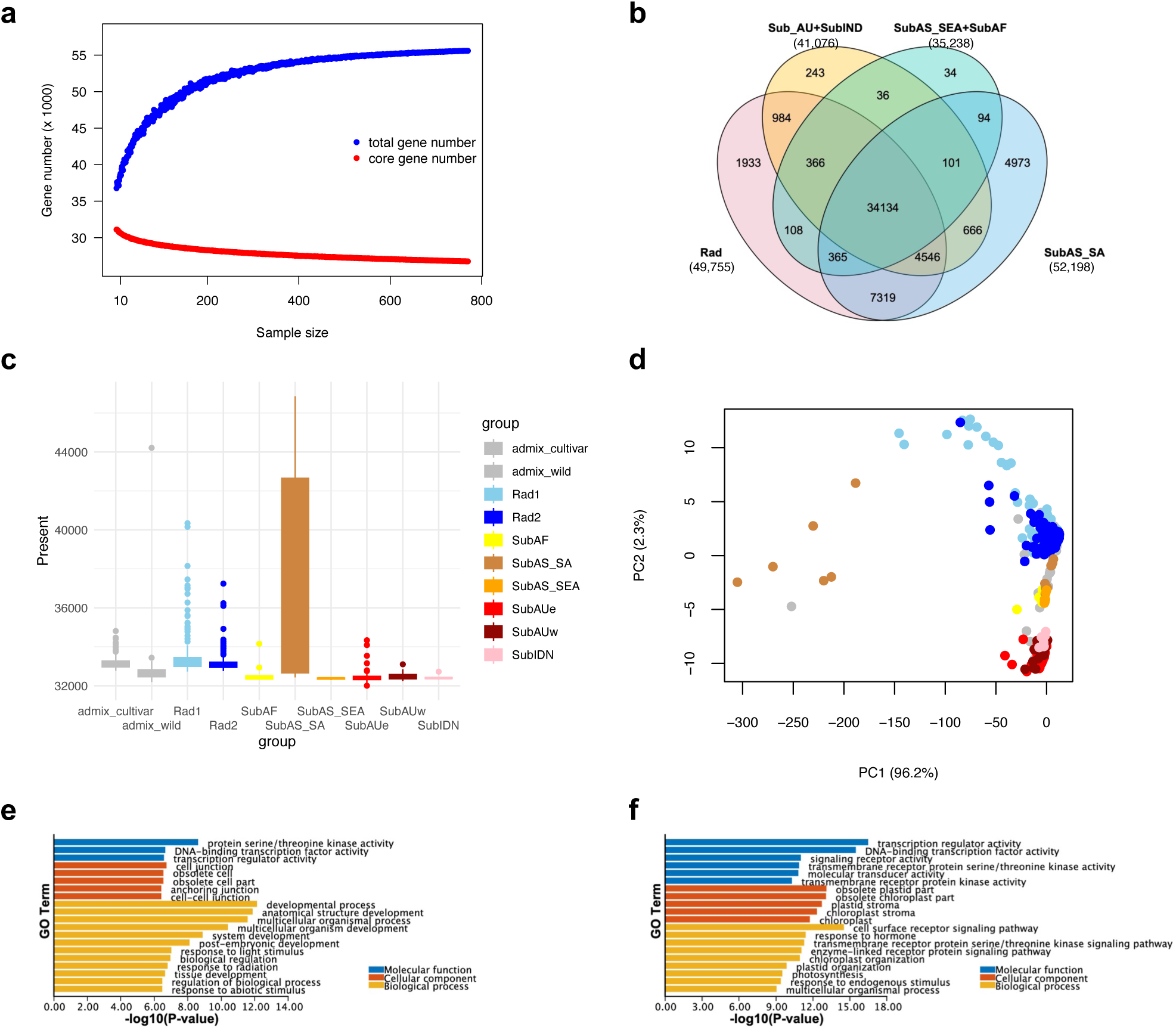
Species-wide pan-genome of mungbean. **a,** Saturation curves showing the total (blue) and core (red) gene counts as a function of sample size. The number of total genes does not plateau until after more than 200-300 accessions were considered, showing the comprehensive amounts of genetic variation in this study. **b,** Venn diagram illustrating the number of genes unique and shared among the genetic groups. **c,** Distribution of gene presences across each genetic group. The box denotes the distribution quantiles, and the whiskers denote 1.5 × interquartile range. Data points beyond the whiskers were plotted individually and considered as outliers. **d,** PCA based on the gene presence-absence variation (PAV) data. **e, f,** Gene Ontology (GO) enrichment analyses in *V. radiata* var. *sublobata* (**e**) and *V. radiata* var. *radiata* (**f**) compared with all genes of the species.

Consistent with their high SNP variation, South Asian *sublobata* have the most genes, with ∼5k unique genes (Fig. 4b). Despite the low SNP variation, cultivars have more unique genes than the other *sublobata* groups (Australian + Indonesian and Southeast Asian + African, grouped based on their phylogenetic relationship). Since SNP variation (mean pairwise nucleotide difference π) is less sensitive to low-frequency variants than gene presence-absence variation (PAV) is, the results suggest cultivars might possess more low-frequency gene presence, and cultivars’ much higher sample size likely enhanced this pattern. The pattern might also result from cultivars being domesticated from another now-extinct Indian *sublobata* group or the introgression practices during mungbean breeding. Consistent with the whole-population pattern, the number of genes in each accession is also higher for South Asian *sublobata*, followed by cultivars (Fig. 4c).

Similar to the patterns of shared SNPs, the cultivars also have more uniquely shared genes (those only exist in cultivars and one specific *sublobata* group) with South Asian (7319 genes, 14% of 52,198 genes in South Asian *sublobata* Fig. 4b) than with other *sublobata* groups (Australian + Indonesian: 984, 2.4% of 41,076, Southeast Asian + African: 108, 0.3% of 35,238), supporting the South Asian domestication scenario. Comparing the PCA from SNPs (Fig. 1b) and PAV (Fig. 4d), some South Asian wild accessions have similar PAV patterns to African wild, Australian wild, and the cultivars, and the PAV neighbor-joining tree showed similar patterns (Supplementary Fig. 15). Given that no obvious gene flow was detected between the cultivar and South Asian wild groups (Supplementary Table 6), some South Asian wild accessions likely still retain the gene PAV patterns close to the original ancestral population where domestication happened.

Compared to the species-wide pan-genome, genes in wild accessions were enriched for gene ontology (GO) functions associated with plant development and responses to light and abiotic stimuli (Fig. 4e), potentially contributing to the adaptive potential to environmental variations across diverse habitats. Meanwhile, the cultivars showed enrichment of genes involved in hormone-mediated regulation and photosynthesis (Fig. 4f), which might be associated with efficient energy accumulation and growth to maximize agricultural productivity in controlled environments.

### Identification of the major-effect QTL across environments

The International Mungbean Improvement Network (IMIN)’s large-scale field trials (Supplementary Fig. 16) allowed us to assess the trait stability and genetic architecture across environments. We first focused on four yield components (days to flowering, pods per plant, seeds per pod, and seed weight) and garden trials with kinship heritability (proportion variation explained, PVE) > 0.2. Seed weight exhibits the highest correlation across trials (Supplementary Fig. 17-20) and demonstrates the highest heritability, reaching a maximum of 0.82 in the field environments.

Performing PAV GWAS separately for each trial, we identified two genes located on chromosomes 11 that were consistently detected across multiple trials (-log_10_*P* > 3 across more than ten trials, Fig. 5a), and they were also detected under GWAS on mean seed weight (Fig. 5b). Notably, Chr11.1340, encodes gamma-glutamyl transpeptidase, whose absence was associated with an increase in seed weight (Fig. 5c). Using the mean seed weight across trials, SNP GWAS of 913630 SNPs revealed nine significant peaks (Fig. 5d; Supplementary Table 9), five of which co-localized with GWAS signals of PAV, overlapping with six significant associated PAV genes (Supplementary Table 10). Interestingly, four of the six significant PAVs, Chr1.2667, Chr1.3000, Chr2.2867, and Chr9.1538, which co-localize with signatures of selection in the landraces (Supplementary Fig. 13), are predominantly present in the wild groups. The higher frequency of gene absence in cultivars (Supplementary Table 11) is associated with larger seeds (Fig. 5e).

**Fig. 5.**
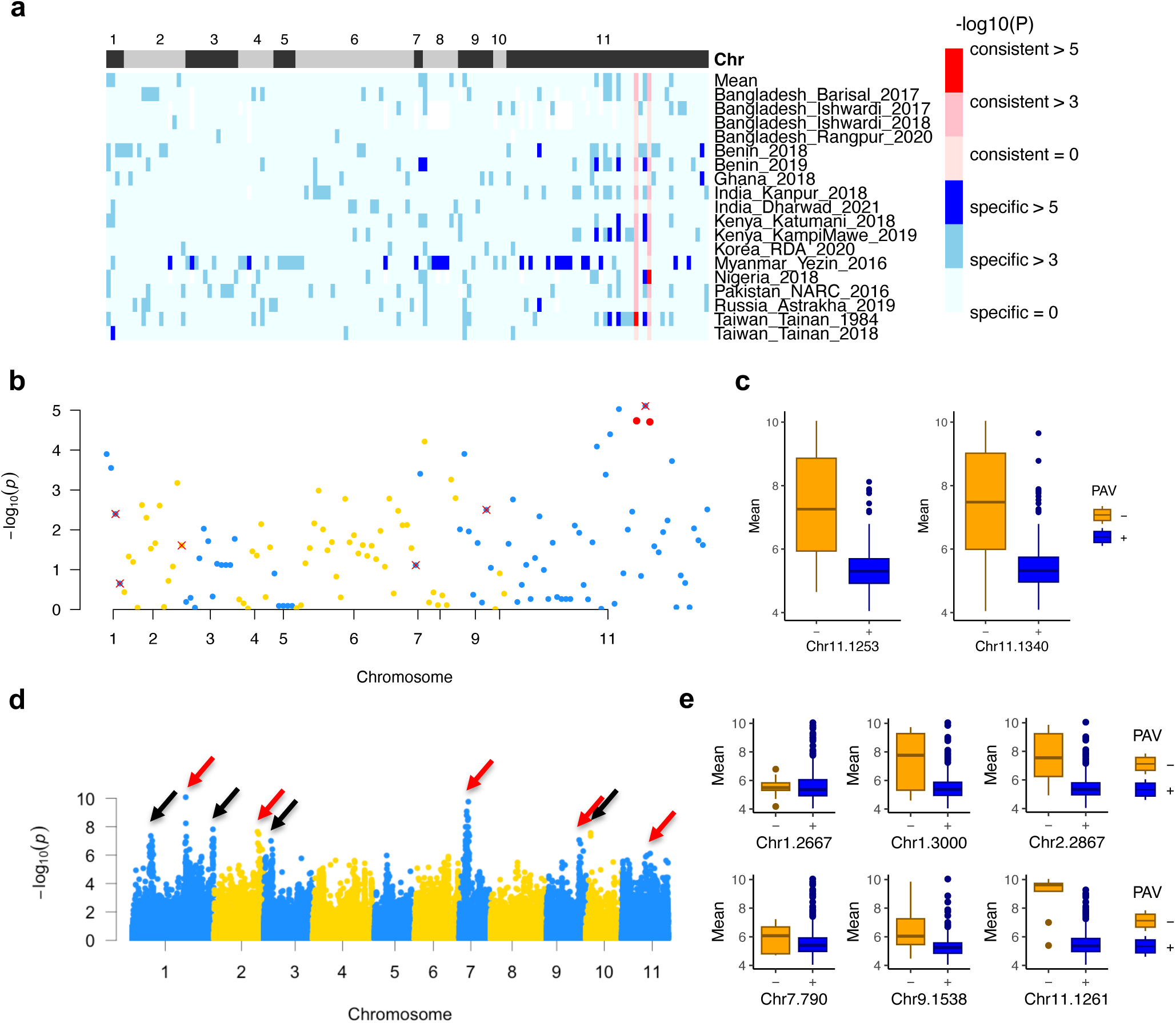
Genome-wide association study (GWAS) of seed weight across multiple environments. **a,** Genes significantly associated with seed weight in the PAV-GWAS across 12 trial sites and multiple years (Bangladesh, Benin, Ghana, India, Kenya, Korea, Myanmar, Nigeria, Pakistan, Russia, Taiwan, and Tanzania). For each gene, color intensity denotes GWAS score (-log_10_*P*) in each trial site, and the red and blue hues denote whether a gene was significantly associated with seed weight across multiple trial sites. **b,** Manhattan plot of PAV-GWAS on mean seed weight. Only genes with log_10_*P* > 3 at any site (those in **a**) were shown. Red dots represent the genes (Chr11.1253 and Chr11.1340) associated with seed weight across multiple trials, and the red x marks indicate genes colocalizing with SNP-GWAS peaks. **c,** Allelic effects of the two genes (those red dots in **b**). **d,** Identification of nine peaks (arrows) based on SNP-GWAS of mean seed weight across trial sites; red arrows indicate loci overlapped with gene presence-absence variation (PAV) GWAS in **a**. To present the potentially polygenic nature of these traits, no hard GWAS threshold was marked. **e,** Allelic effects of six present/absent genes (red arrows in **d**) colocalizing with SNP-GWAS loci.

### Genomic prediction for native environments and transcontinental GxE interaction

IMIN’s trans-continental trials allowed the identification of stable traits across environments (seed size, Fig. 5) and facilitated our dissection of gene-environment associations (Supplementary Fig. 16). We treated collection-site environmental variables as traits and used 178 accessions with GPS coordinates to predict the native climatic conditions of 1092 landraces across Asia^12^. A total of 34 environmental variables with high environment-genome association (PVE > 0.5, Supplementary Table 12) were used in five genomic prediction methods (rrBLUP, GBLUP, RF, SVM, and BSLMM). The accuracy and error vary across environmental variables and methods (Supplementary Fig. 21 & Supplementary Table 13), and for the following analyses, predictions from RF were used due to its generally better performance.

The 1092 landrace accessions were separated into four genetic groups from South Asia, Southeast Asia, East Asia, and Central Asia^12^. The predicted climate of origin is consistent with their genomic background. For example, accessions belonging to the southern groups were predicted to have lower latitudinal distribution and higher summer growing-season precipitation than northern groups (Fig. 6ab). PCA of 17 environmental variables (r < 0.95, Supplementary Fig. 22, Supplementary Table 14) recapitulates accession geographical distribution and the IMIN field environments (Fig. 6c and Supplementary Fig. 23), with PC1 explaining almost 80% of the total variation, consistent with the previous results that the main environmental and phenotypic difference exists between the southern and northern Asia^12^.

**Fig. 6.**
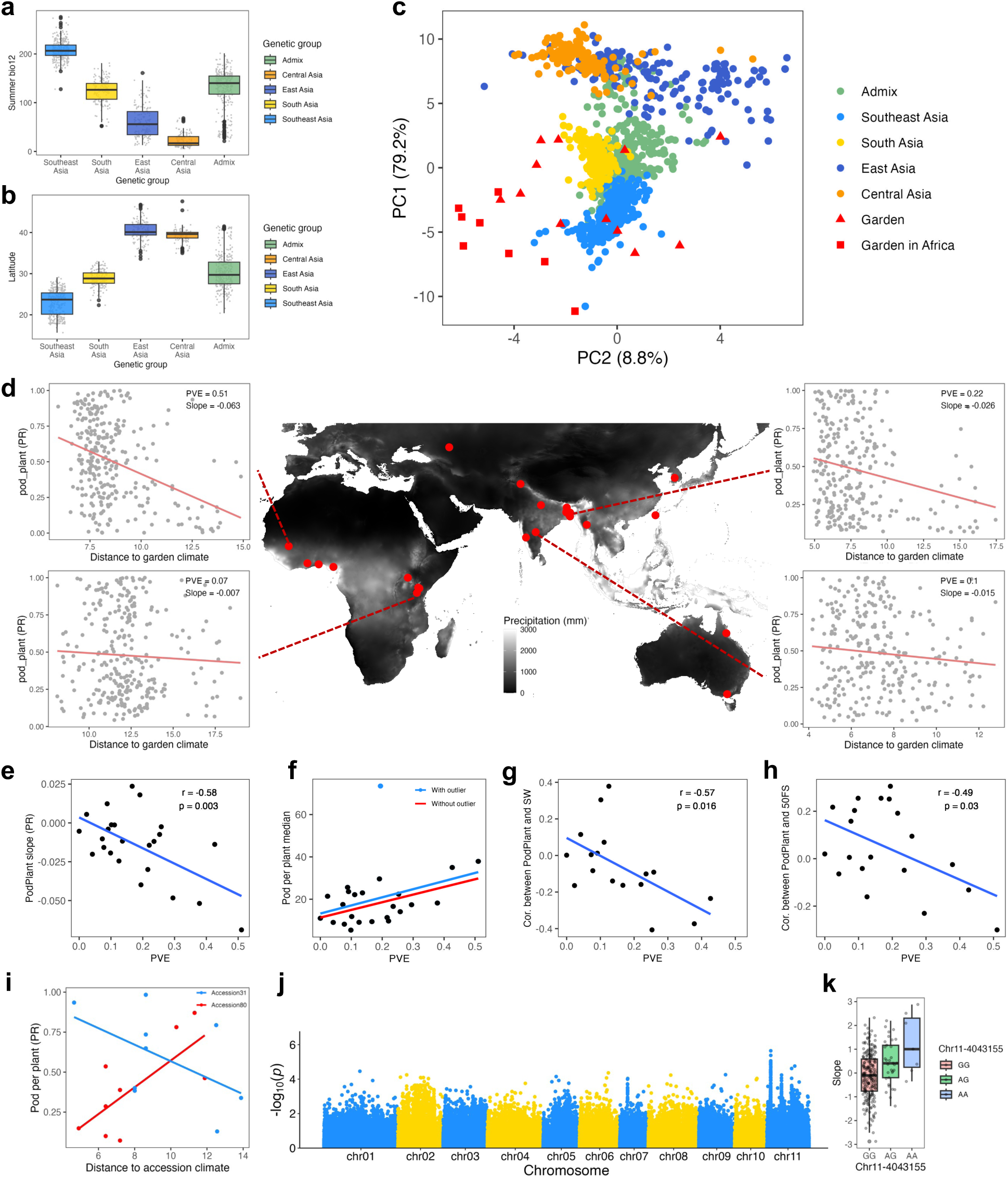
Native environment prediction and factors affecting mungbean yield across multi-environment trials. **a-b,** Boxplots showing native environment of accessions in major cultivar genetic groups, estimated by genomic prediction. Two exemplary variables (summer precipitation bio12 and latitude) are shown. **c,** Principal component analysis (PCA) of predicted environmental variables reveals clear separation among genetic groups. **d,** The trial sites of the International Mungbean Improvement Network. Number of pods per plant is negatively associated with the climatic distance of each accession’s native site to the trial site, suggesting that accessions perform better under environments similar to their predicted origin. **e,** The extent of local adaptation (regression slow in **d**) is highly associated with genome-level heritability (PVE) of the trait in these sites. **f,** Sites with higher yield heritability (PVE) also have higher overall yield. Two regression lines are shown, including (blue) or excluding (red) an outlier site. **g,** Sites with higher yield heritability (PVE) have a more negative correlation between pod number and seed weight. **h,** Sites with higher heritability have a more negative correlation between pod number and flowering speed. **i,** Reaction norms of two exemplary mungbean accessions across trial sites. The horizontal axis denotes the distance of an accession’s native environment to the trial-site environment. The blue accession (with a negative slope) shows signs of local adaptation. **j,** Genome-wide association study (GWAS) results using the reaction norm slope as a trait. To present the potentially polygenic nature of these traits, no hard GWAS threshold was marked. **k,** Boxplot showing allelic effects on reaction norm slopes for Chr11–40315155.

Combining the predicted native environments and the field data of IMIN trials, we investigated factors affecting the major yield component, pods per plant. If local adaptation exists in landraces, well-performing accessions should be those whose native environments are similar to the cultivation site, i.e., a negative association between performance and the origin-garden environmental difference. The pattern, however, varies across sites and years (Fig. 6d and Supplementary Fig. 24). Among sites, kinship heritability (PVE) has a strong negative correlation with the performance-environmental difference regression slope (calculated separately for each garden, Supplementary Fig. 24), suggesting the pattern of local adaptation is more pronounced in a growing condition with high heritability, i.e. manifesting the genetic difference among accessions (Fig. 6e). Adjusting the threshold of including less-correlated variables to calculate environmental distance does not change this conclusion (Supplementary Fig. 25).

The pattern could be explained by either the stress tolerance or fecundity hypotheses (Supplementary Fig. 26). The former predicts higher PVE in low-yield conditions (allowing tolerant accessions to thrive), and the latter predicts the opposite. Except for Hyderabad, India (the outlier in Fig. 6f), there is a strong positive relationship between overall performance and the PVE within gardens (Fig. 6f), supporting that benign environments (or good agricultural management) increase population performance by enabling the potential of high-yield accessions, thereby increasing the PVE and exhibiting local adaptation.

While an ideal accession should have abundant pods per plant, large seeds, and fast flowering, tradeoffs among yield components exist. Interestingly, the magnitudes of tradeoffs (between-trait correlations) differ among gardens, and stronger tradeoffs (negative correlations) between pod per plant and seed weight (Fig. 6g) or flowering speed (50FS, the negative values of days to 50% flowering, Fig. 6h) exist in gardens with larger pod-per-plant PVE. The results imply that while growing conditions maximizing the genetic effect on yields (high PVE) might facilitate selective breeding and enable the potential of accessions from similar climatic conditions, such conditions also strengthen the tradeoffs among yield components, constraining the efficacy of selection.

Using similar approaches but focusing on each accession’s performance across gardens, we investigated the reaction norms reflecting accession plasticity across environments. Here, a negative correlation between accession performance and its native-site environmental distance to different gardens suggests local adaptation (Fig. 6i). By treating such a slope as a quantitative function-valued trait, we performed GWAS to identify plasticity-altering loci. We identified a peak on chromosome 11 (Fig. 6j, the red arrow), with the rare allele increasing the slope, exhibiting a lack of local adaptation and is likely selected against (Fig. 6k). This region spans approximately 500 kb and contains multiple candidate genes, most of which are related to proteins involved in ubiquitination, likely playing regulatory roles in protein signaling for mungbean adaptation to various environmental conditions^23^.

## Discussion

While many works have investigated the genomic variation of crops, it is essential to note that the research scope heavily depends on the sampling design. This work compiled the most comprehensive genomic variation of mungbean, a crop important for worldwide small-holder farmers. Such a collection enables the investigation of species-wide variation, facilitates the dissection of demographic history, unveils a unique case of parallel phenotypic evolution by parallel mutations of the same locus, allows the identification of more pan-genes, and suggests novel strategies for future breeding programs accounting for genotype-by-environment interactions.

Incompatible with archaeological evidence that mungbean was domesticated in South Asia^33^, the nuclear genome showed that the South Asian wild group is the most divergent from all other groups (Fig. 1). On the other hand, some Indian wild accessions possess chloroplast haplotypes closest to those of cultivars (Fig. 2a). The cultivar and South Asian wild groups also shared more SNPs (Supplementary Fig. 12) and pan-genes (Fig. 4b), and some South Asian wild accessions have similar gene PAV patterns as the cultivars (Fig. 4d). These results suggest that the wild group closest to the direct progenitors of modern cultivars might have been extinct or were not sampled in this work, while residual traces of similarity could still be detected in chloroplast haplotypes or parts of the nuclear genome. This might be similar to rice, where no current wild populations exist near the Yangtze River Basin^34^. Since crops might require habitats similar to those of their wild progenitors, it has been suggested that the expansion of crop cultivation may be associated with the loss of wild populations^35^.

Unlike previous mungbean studies focusing on regional collections^15^, this work incorporates wild populations across the species range and cultivars spanning global climatic gradients. The comprehensive collection allows the identification of more pan-genes, which do not saturate until >400 accessions were considered (Fig. 4). The pan-genome resource and multi-environment trials also allow us to identify candidate gene PAVs stably affecting seed size across environments (Fig. 5). Uniquely incorporating the prediction of accession native environments and large-scale field trials, we further showed that local adaptation exists when a trial has high heritability, and the high heritability was manifested by a benign environment/management increasing overall yield, facilitating the potential of high-yield accessions (Fig. 6). For future breeding targets, our results suggest one might focus on selecting high-yield accessions in benign environments, and these accessions most likely originated from a similar environment to the trial sites (Fig. 6).

By incorporating the most comprehensive species-wide genomic variation and field trials, we unveiled the complex demographic history and used this to inform future breeding and cultivation strategies.

## Methods

### Plant materials and sequencing

A worldwide collection, including 618 *V. radiata* var. *radiata* accessions, 162 *V. radiata* var. *sublobata,* and four outgroup accessions, *V. mungo*, were whole-genome re-sequenced. The passport information can be found in Supplementary Table 2. For the *V. radiata* var. *radiata* materials, the genomic DNA was extracted from leaves using FavorPrepTM Plant Genomic DNA Extraction Kit (FAVORGEN). 40X coverage of whole genome re-sequencing was provided by the Twelfth Annual Illumina Agricultural Greater Good Initiative Grant. For the *V. radiata* var. *sublobata* materials, the genomic DNA was extracted from leaves with the DNeasy Plant Mini Kit (QIAGEN). DNA libraries were constructed using NEBNext Ultra™ II DNA Library Prep Kit, and 20X coverage was completed using Illumina HiSeq X Ten.

### Nanopore sequencing

One cultivated commercial accession (Crystal) and two wild accessions from Australia (CQ3649) and India (Karumbyar) were chosen. High molecular weight genomic DNA (gDNA) was extracted following the protocol for long-read sequencing^36^. Briefly, gDNA of 3.5 g of frozen fresh leaves was extracted with the lysis buffer (1% Polyvinylpyrrolidone 40, 1% Sodium metabisulfite, 0.5 M Sodium chloride, 100 mM Tris, 50 mM EDTA, 1.25 % SDS). Then, protein residues were removed via a phenol-chloroform-isoamyl alcohol mixture (25:24:1). Finally, gDNA was captured with AMPure XP beads (Beckman Coulter) and eluted with TE buffer. Before DNA library construction, the gDNA fragments were depleted using Short Read Eliminator (SS-100-101-01, Circulomics).

The 1D-ligation library prep kit (SQK-LSK109, Oxford Nanopore Technologies) was used for Nanopore library construction, and the library was sequenced using MinION flow cells (version R9.4). The raw sequencing reads were called with Guppy v3.2.2 (Oxford Nanopore Technologies).

### Genome assembly

Raw Nanopore reads were trimmed by Porechop v0.2.4^37^ to remove adaptor sequences and then assembled using Flye v2.5^38^ with a read length over 10 kb. The consensus sequences of the draft assemblies from three accessions by Flye were polished using Illumina reads using Pilon v1.23^39^ with three iterations. The genome size of each accession was estimated using Jellyfish v2.2.10 on the Illumina short reads to obtain the 25-base k-mer frequency distribution and following the genome size estimation tutorial on the website (https://bioinformatics.uconn.edu/genome-size-estimation-tutorial/#). The completeness of assemblies was evaluated using Benchmarking Universal Single-Copy Orthologs (BUSCO) v4.0.6^40^ with an embryophyta dataset of 1,614 plant-specific orthologous genes (lineage dataset embryophyta_odb10).

To further refine the Crystal and CQ3649 accession genomes, we anchored chromosome-scale sequences with linkage map^41^. Recombinant inbred lines from two mungbean crosses, V2802 x NM94^4^ and Dahuaye x Jilyu 9-1^42^ were used to construct linkage maps with MSTmap v1^43^. The following parameters were used in MSTmap: distance_function kosambi, no_map_dist 15.0, no_map_size 2, missing_threshold 0.25, estimation_before_clustering yes, detect_bad_data yes, and objective_function ML. The cut-off p-value was determined when markers were assigned to 11 linkage groups. ALLMAPS v1.0.9^44^ combined the two linkage maps and further scaffolding draft assembly into chromosome-level. For Karumbyar accession, Ragtag^45^ was used to construct a complete genome using the Crystal genome as a reference based on the homology-based scaffolding method.

To remove scaffolds belonging to the organellar genome, the unanchored scaffolds were aligned to the mungbean mitochondrial and chloroplast genomes using minimap2 v2.2^46^. The aligned scaffolds were discarded from the nuclear genome assembly.

### Gene prediction and annotation

We implemented *ab initio*-based, transcriptome-based, and homology-based gene structure prediction strategies to identify protein-coding genes. For the *ab initio* gene prediction, AUGUSTUS v3.3.3^47^ was used with the -species = arabidopsis parameter. For the transcriptome-based gene prediction, we used two types of evidence: transcript assembly and predicted open reading frames (ORFs). Transcriptomic data were obtained from our collection as well as the public database. RNA-seq data were extracted from distinct mungbean tissues. Public RNA-seq was downloaded from the NCBI database in SRA archives SRP043316, SRR1407784, and BioProject PRJNA276314. Those RNA-seq data were mapped to the Crystal genome using HISAT2 v2.2.0 and aligned to generate transcriptome assembly by StringTie v2.1.2^48,49^. In addition to reconstructing transcripts, we also identified candidate ORFs using TransDecoder v5.5.0 (https://github.com/TransDecoder) for those longer than 100 amino acids by integrating the UniProt and Pfam domains search results into coding region selection. For homology-based gene prediction, protein sequences of *Arabidopsis thaliana* (GCF_000001735.4), *Glycine max* (GCF_000004515.6), *Vigna angularis* (GCF_001190045.1), *V. radiata* (GCF_000741045.1), and *V. unguiculata* (GCF_004118075.1) were downloaded from NCBI database. These protein sequences from these plant species were aligned to the reference genome by GenomeThreader v1.7.1^50^. All predicted results were integrated into EVidenceModeler v1.1.1 (EVM)^51^ with the evidence weight sets, AUGUSTUS: 1, StringTie: 10, TransDecoder: 5, GenomeThreader: 5, to generate weighted consensus gene structures.

To obtain functional annotation of each annotated protein-coding gene, we searched them against different public databases using multiple tools, including KAAS^52^, BLASTP v2.9.0+^53^, eggNOG-mapper v2.1.9^54^, Mercator4 v5.0^55^, and TRAPID^56^. For KAAS, transcript sequences were mapped to the KEGG ortholog database^57^, representing the molecular function of orthologs. For BLASTP, the BLASTP v2.9.0+ algorithm was used for searching homologous proteins by comparing our protein sequences to the Swiss-Prot/plant taxonomy (https://www.uniprot.org/, accessed: 2022-08-07) and NR protein databases (NCBI non-redundant protein sequences, accessed: 2022-08-08). Based on the sequence similarity, the best match of the alignment was used to describe the protein’s function. The eggNOG-mapper derived orthologous groups from the eggNOG v5.0 database^58^, which includes the functional annotation of eggNOG ortholog groups (OGs), KEGG, and GO (Gene Ontology)^58^. In addition, we used the online plant protein annotation tool Mercator4 to annotate the plant biological functions of the protein. TRAPID, which includes several databases such as Monocots PLAZA-4.5.1^59^ and Pico PLAZA-3.0^60^, was also used to assess GO and IPR (Inter-Pro) mappings.

### Organelle de novo assembly and gene annotation

Unicycler^61^, which combined long and short-read data, was used to assemble organellar genomes. The raw reads belonging to organellar assembly were baited using reference published mungbean organellar genomes downloaded from NCBI (NC_015121.1 for mitochondrion genome and NC_013843.1 for chloroplast genome). Raw Nanopore long and Illumina short reads were mapped to the reference organellar genomes using minimap2 v2.2^46^ and BWA v0.7.15^62^, respectively. Mapped reads were extracted using bedtools^63^. Reads longer than 5 kb and read quality score over nine were retained using Nanofilt^64^ for the following organellar assembly. Since Crystal’s chloroplast genome could not be assembled entirely using Unicycler, Getorganelle^65^ with default parameters was used. The Geseq^66^ was used to predict and annotate the genes within the complete organellar genomes.

### Pan-genome construction

After cleaning and remapping the raw Illumina sequencing reads, we used samtools to grep unmapped reads and then used ABySS 2.0 to assemble these unmapped reads^67^. Contigs less than 500 bp were removed. CDHIT was used to remove redundant genes across the pan-genome^68^. In addition, contigs matched to virus, protozoa, fungi, and bacteria databases in NCBI were removed. All the contigs that can be blasted back to plant-specific databases, including InterPro, InterPro_des, Mercator, and Swiss_port, were kept as the final extra genomic contigs. The annotation followed the same procedure as the **Gene prediction and annotation** section. After aligning cleaned reads back to the pan-genome, the presence/absence variants were estimated by SGSGeneLOSS following the manual^69^.

### Variant calling

The SNP calling was following the methods in Lin et al.^22^. In brief, the raw sequencing reads were cleaned with SolexaQA++ v3.1.7.1 and mapped to Crystal reference genome with Burrows-Wheeler aligner v0.7.15^62,70^. Duplicated reads were marked with Picard v2.9.0-1 (http://broadinstitute.github.io/picard/). SNPs were called with GATK v4.1 individually by the HaplotypeCaller and then merged by GenotypeGVCFs^71^. SNPs were filtered by vcftools v0.1.13 with the following parameters “missing rate < 0.1, --min-alleles 2 --max-alleles 2 --remove-indels --max-missing 0.9 --minQ 30”^72^.

### Population genomics analyses

Bi-allelic SNPs were used for population genetics analysis. For linkage disequilibrium (LD) and fixation index (FST) analyses, SNPs with minor allele frequency (MAF) less than 0.05 were further removed. Population structure was inferred, and a suitable number of clusters was evaluated with cross-validation error using ADMIXTURE ^73^. Principal component analysis (PCA) of 780 mungbean accessions was conducted using PLINK v2.0 alpha^74^. Pairwise fixation index (FST) between genetic groups and nucleotide diversity (π) in non-overlapping 10-kb sliding windows were calculated using vcftools v0.1.13^72^. The pairwise linkage disequilibrium (LD) between SNPs was calculated with PopLDdecay v3.41^75^. To detect the ancient introgression between two populations, we ran ABBA-BABA test (D-statistics) and calculated D-values using Dsuite^76^ to provide evidence of ancient introgression. The outgroup f3 statistics were measured using the ADMIXTOOLS^77^ package in R.

### Phylogenetic relationship

SNPs on chromosomes were filtered by SNPhylo v.20180901^78^ with parameters “-m 0.001 -M 0.01 -l 0.5” and generated the aligned sequences. Then, the sequences were used to construct the maximum likelihood tree by FastTree2 v2.1.11^79^ with parameters “-nt –gtr”. Treemix v.1.13^80^ was performed to infer population split and migration models between genetic groups. LD-pruned SNPs were obtained using PLINK v2.00a3.6LM “--indep-pairwise 50 10 0.2”^74^. Then Treemix was run on allele frequencies from an LD-pruned SNP dataset of accessions from eight genetic groups (30 accessions in each group were randomly chosen) with 500 bootstrap replicates to build a population-based maximum likelihood tree, considering migration events to improve the fit of the inferred tree. OptM v0.1.5^81^ was used to estimate the optimal number of migration events on a population-based tree generated by Treemix.

The R package “ape”^82^ was used to construct a neighbor-joining tree of the chloroplast genome of 780 mungbean accessions. The haplotype network based on the chloroplast genome was built using PopART with the medium-joining algorithm^83^.

The SNPs without heterozygotes and missing values were used for the haplotype network analysis with four *V. mungo* accessions as the outgroup.

### Phenotypic evaluation and relationship of data among multiple locations

The mungbean mini-core collection was shared with the partners of the International Mungbean Improvement Network, including the national agricultural institutes in Australia, Bangladesh, Benin, Ghana, India, Mali, Myanmar, Kenya, Nigeria, Pakistan, Russia, Taiwan, Tanzania, and Uganda. The phenotypic evaluation of the mini-core collection was conducted by these partners from 2016 to 2020 in multiple locations (Supplementary Table 15). Days to 50% flowering (DF), average seed weight per pod (seed per pod), average pod number per plant (pod per plant), 100 seed weight (SW), and seed coat color were evaluated. To investigate the relationship of the phenotypic data among multiple locations, we conducted the Principal Component Analysis (PCA) and Pearson correlation analysis using “ade4”^84^ and “stats”^85^ packages in R, respectively. For GWAS, genetic variants were filtered by MAF < 0.05. GWAS were conducted by Genome-wide Efficient Mixed Model Association (GEMMA) with default settings^86^.

### Genomic prediction for native environments and G x E interaction

For the original 1092 worldwide mungbean landraces, only the 178 accessions from the Vavilov Institute have collection-site GPS coordinates. For accessions with GPS coordinates, we extracted climatic variables, including 42 bioclimatic variables from WorldClim version 2^87^, five variables from ENVIREM^88^, four variables from CHELSA^89^, and one from FAO (Supplementary Table 12). Treating these environmental variables as traits, we ran a null model in TASSEL 5.0^90^ to estimate the proportional trait variation explained (PVE) by the genetic kinship matrix. Only those with PVE higher than 0.5, suggesting that the native environments have a strong genetic basis and local adaptation, were used in further analyses. Genomic selection methods, including ridge regression best linear unbiased prediction (rrBLUP^91^), genomic best linear unbiased prediction (GBLUP^92^), random forest (RF^93^), support vector machine (SVM^94^, and Bayesian sparse linear mixed model (BSLMM^95^) were used to predict the collection-site environments for those without original GPS coordinates. Each method was repeated 100 times, among which the RF and SVM methods use 5-fold cross-validation. The final model fit was judged by Pearson’s correlation coefficient *r* between observed and predicted values in the training sets; only the climatic variables with *r* > 0.7 were used for the following analyses. Mean absolute error (MAE) and mean square error (MSE) were also calculated to ensure the difference between observed and predicted values was minimal. Pairwise *r* was further calculated for the predicted values of the 30 remaining environmental variables, and we only kept 17 variables whose mutual *r* was less than 0.95 for the following analyses (Supplementary Table 14).

Based on the predicted or observed values of 17 environmental variables, we calculated the Euclidean distances between all accessions and all gardens. For the 292 WorldVeg Minicore accessions with field trial data across multiple gardens, we investigated the relationship between their yield (pods per plant) and the absolute value of accession-garden environmental distances. Focusing on all accessions’ yield in a given garden, this analysis investigated whether a garden’s environment facilitated the exhibition of local adaptation, where negative slopes represent higher yield for accessions whose native environment was closer to this garden. Focusing on one accession’s yield (transformed to rank percentile within each environment, removing the garden main effect) across gardens, this analysis investigated whether an accession exhibited the property of local adaptation, where negative slopes represent higher rank yield in gardens with environments closer to its native site. For both analyses, the differences in slopes among gardens or accessions represent G x E interaction. Finally, to investigate potential genetic bases of such G x E response, we performed GWAS for the “slope across garden” calculated for each accession.

### Seed color genes, constructs, and transgenic plants

The seed coat color of MMC accessions was recorded in Mareeba, Australia, in 2016 with five categories (1: yellow, 2: green, 3: brown, 4: speckled, 5: black). Since few accessions belong to category 1 and no clear distinction exists among category 3-5, we coded the phenotype as green vs. dark (category 3-5) and used this binary trait for GWAS. GEMMA (version 0.98.3)^86^ was used to calculate population kinship matrices for the filtered SNP datasets (max missing 0.1, MAF 0.05). GWAS was performed using a univariate mixed linear model method (LMM) with the default parameters in GEMMA. The Manhattan plot was generated using the R library qqman package^96^.

The Coding DNA Sequence (CDS) of the wild-type *VrPAP1* was cloned from the Karumbyar accession as the genomic background using PCR primers (F: 5’ATGGAAGAATTATCAGGCGTGAG and R: 5’CTAATTAATTTCTAGGTCAAGAAACATATCGC). The resulting PCR fragment was inserted into the TA-vector, pCR8/TOPO/TA (ThermoFisher Scientific).

Subsequently, the *VrPAP1* CDS region was transferred into pEarleyGate 100 destination vector^97^, which contains a constitutive 35S promoter, using Gateway LR Clonase II (ThermoFisher Scientific). The *VrPAP1* expression construct was introduced into *Agrobacterium tumefaciens* strain GV3101, which was then used to transform *A. thaliana* via the floral-dip method. For *A. thaliana*, the homolog mutant *Attt2-5* (CS105593), characterized by a lighter seed coat phenotype than its wild progenitor, was used as the genomic background for this assay. Homozygous T₃ seeds overexpressing *VrPAP1* were selected for seed coat color analysis. Siliques were harvested one and two weeks after flowering, and seeds were observed and photographed under a stereomicroscope. Additionally, mature dry seeds were collected and imaged using microscopy for further phenotypic characterization.

## Acknowledgement

We thank the Illumina Agricultural Greater Good Initiative Grant for supporting sequencing of mungbean cultivars and the IMIN partners for conducting field works. We thank Dr. Kingsley Uzoma (Michael Okpara University of Agriculture, Nigeria), Dr. K. Offei Bonsu (CSIR-Crops Research Institute, Kumasi, Ghana), and Dr. Cho Gyu Taek (National Institute of Crop and Food Science, Republic of Korea) for their help. We are grateful to the support from Computer and Information Networking Center, National Taiwan University for the high-performance computing facilities as well as Technology Commons, College of Life Science, National Taiwan University for the molecular biology assistance. C-RL was supported by 111-2628-B-002-021 and 112-2628-B-002-023-MY3 from National Science and Technology Council, Taiwan. RS and RMN were funded by the Australian Center for International Agricultural Research (ACIAR) through the projects on International Mungbean Improvement Network (CIM-2014-079 and CROP-2019-144) and by the strategic long-term donors to the World Vegetable Center: Republic of China (Taiwan), United States Agency for International Development (USAID), the UK Government’s Foreign, Commonwealth & Development Office (FCDO), ACIAR, Germany, Thailand, Philippines, Korea, and Japan. MGS was supported by the SPbPU Fund for the Advancement of Science and Technology.

## Data availability

All sequence and assembly data are available at NCBI BioProject PRJNA1338092. Trait data and related codes were deposited at https://github.com/YaPing-omics/Global-Mungbean-PanGenome.

