## Supplement Figure for "The demographic history, genomic variation, and transcontinental genotype-phenotype-environment map of mungbean"

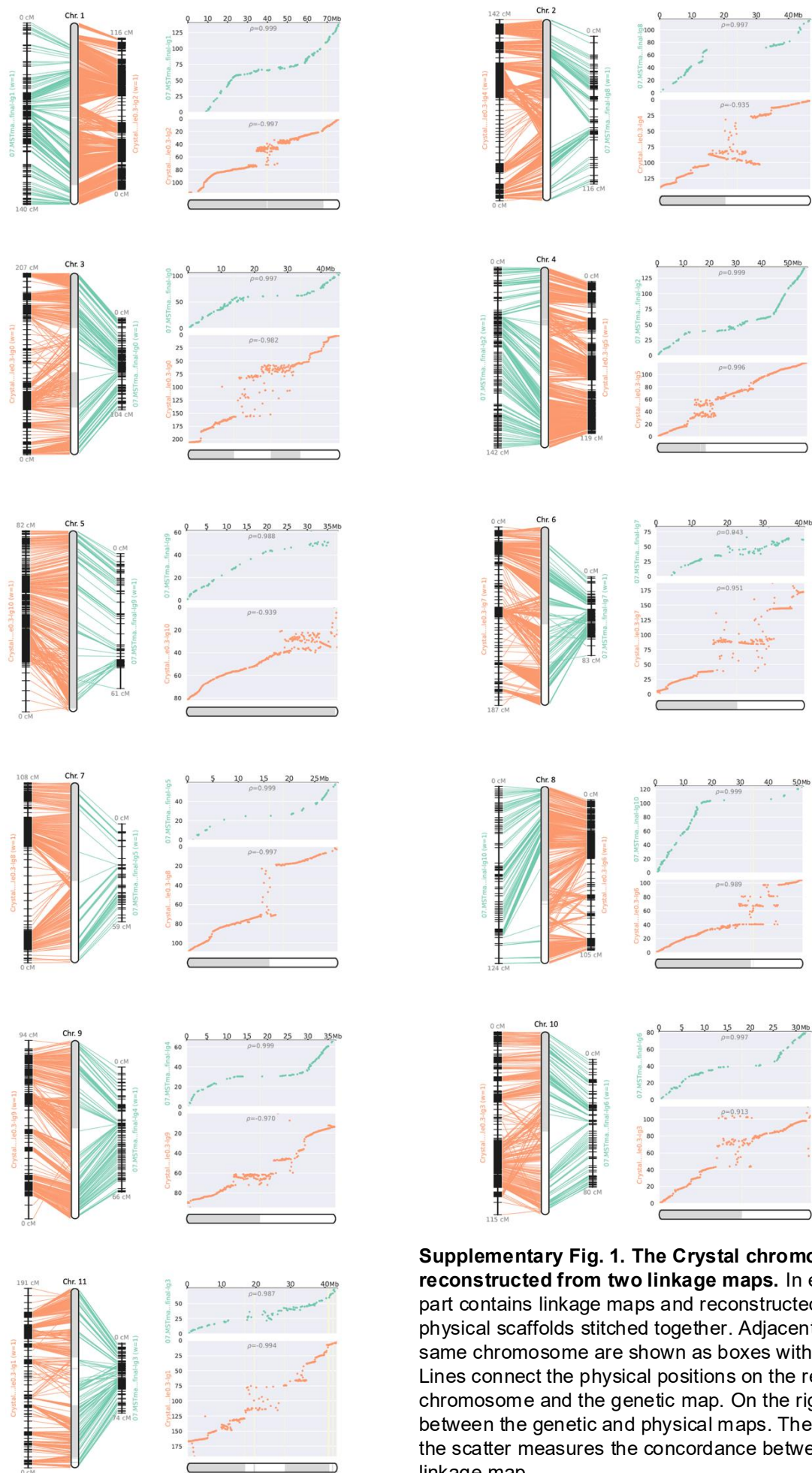

**Supplementary Fig. 1. The Crystal chromosome reconstructed from two linkage maps.** In each panel, the left part contains linkage maps and reconstructed chromosomes with physical scaffolds stitched together. Adjacent contigs within the same chromosome are shown as boxes with alternating colors. Lines connect the physical positions on the reconstructed chromosome and the genetic map. On the right are scatter plots between the genetic and physical maps. The Spearman's $\rho$ on the scatter measures the concordance between the physical and linkage map.

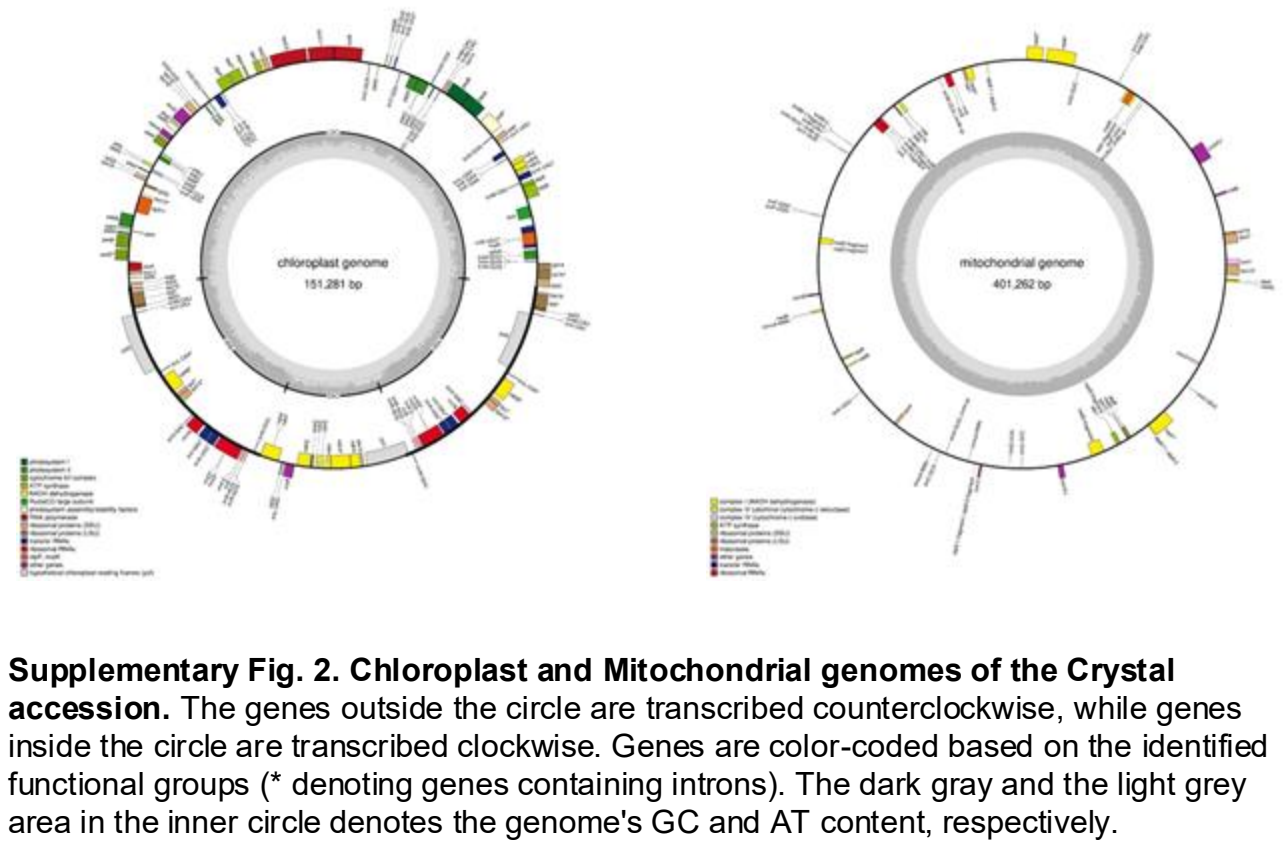

Supplementary Fig. 2. Chloroplast and Mitochondrial genomes of the Crystal accession. The genes outside the circle are transcribed counterclockwise, while genes inside the circle are transcribed clockwise. Genes are color-coded based on the identified functional groups (\* denoting genes containing introns). The dark gray and the light grey area in the inner circle denotes the genome's GC and AT content, respectively.

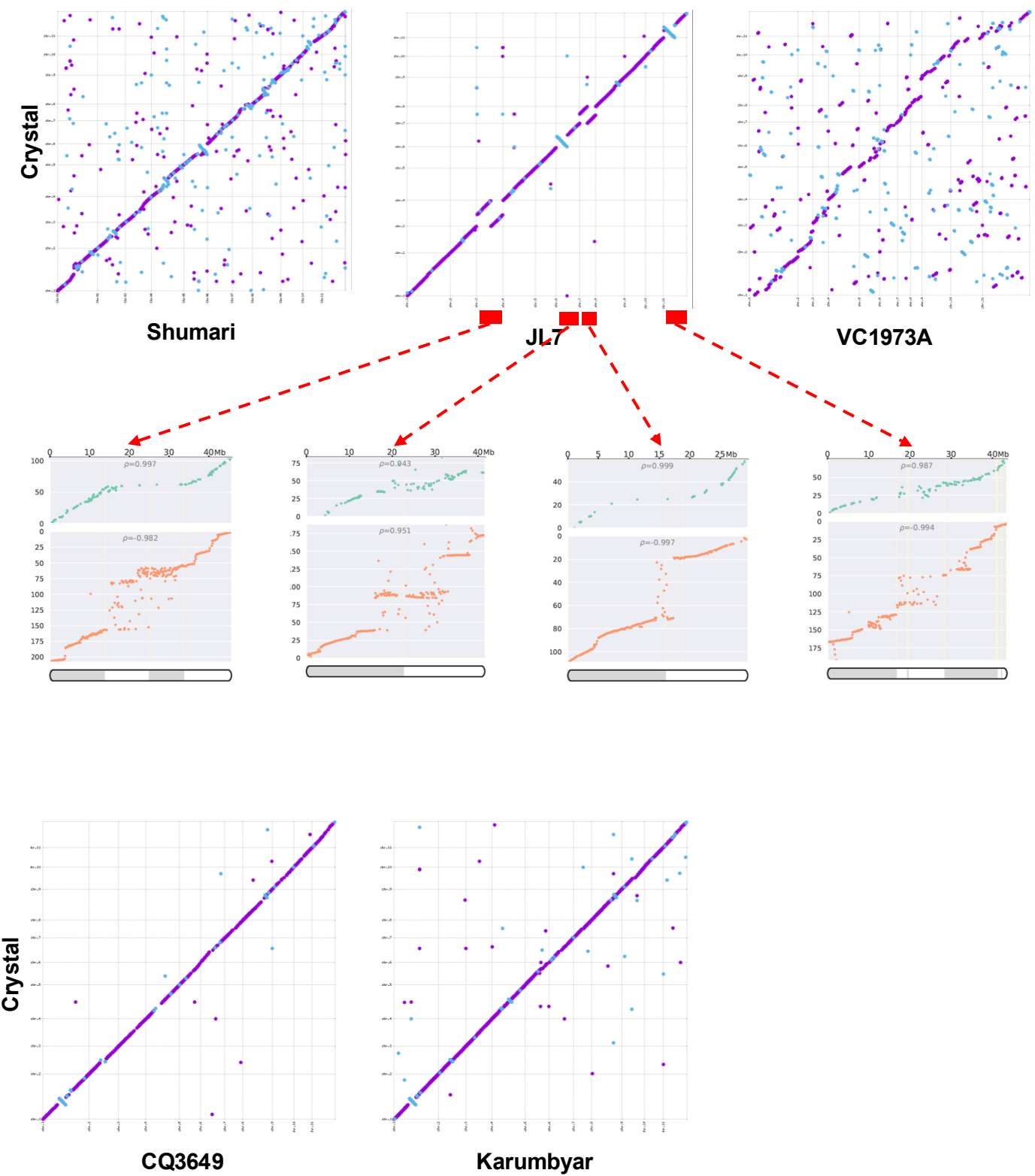

**Supplementary Fig. 3. MUMmerplot comparison of Crystal and other genomes (*V. angularis* - Shumari, *V. radidata* – JL7, VC1973A, CQ3649, and Karumbyar).** Chromosomal translocations and inversions were observed between Crystal and JL7, and for these chromosomes, the linkage-physical map correspondence was shown for Crystal.

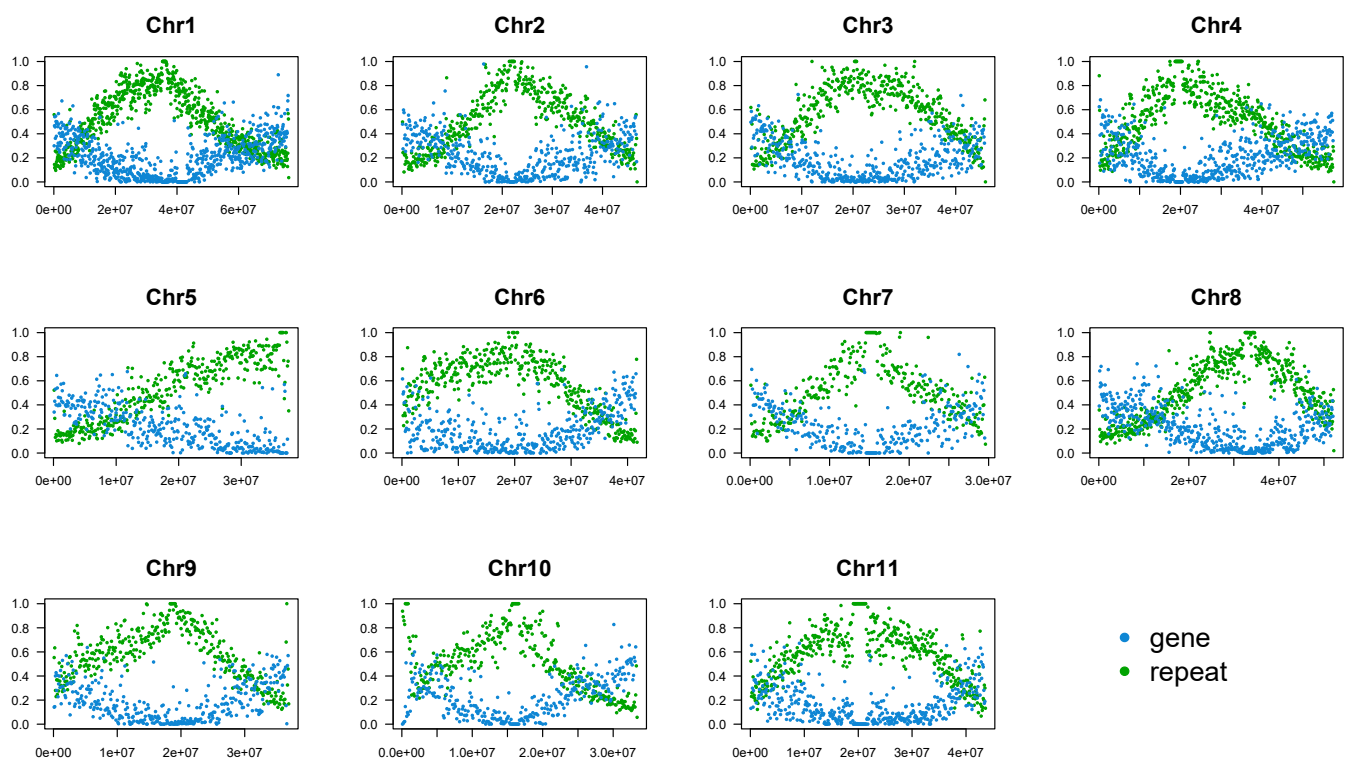

**Supplementary Fig. 4. Gene and repeat density across the Crystal genome.** The proportions of sequences belonging to annotated genes or repetitive regions within each 100kb window are shown.

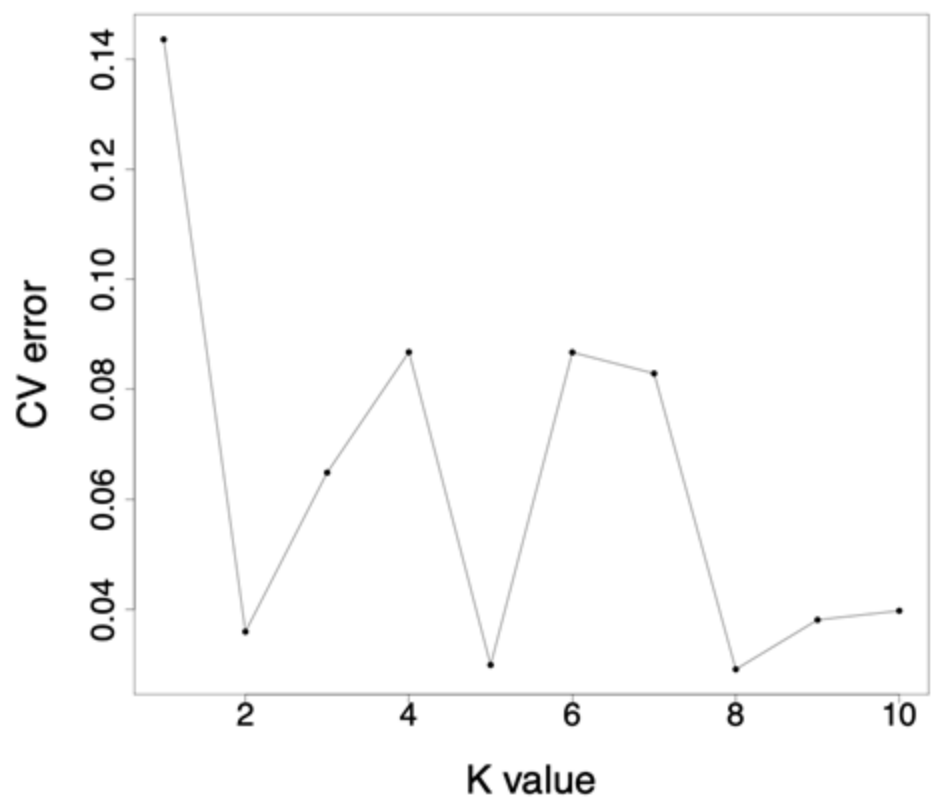

**Supplementary Fig. 5. Cross-validation error of each K value in the ADMIXTURE analysis.**

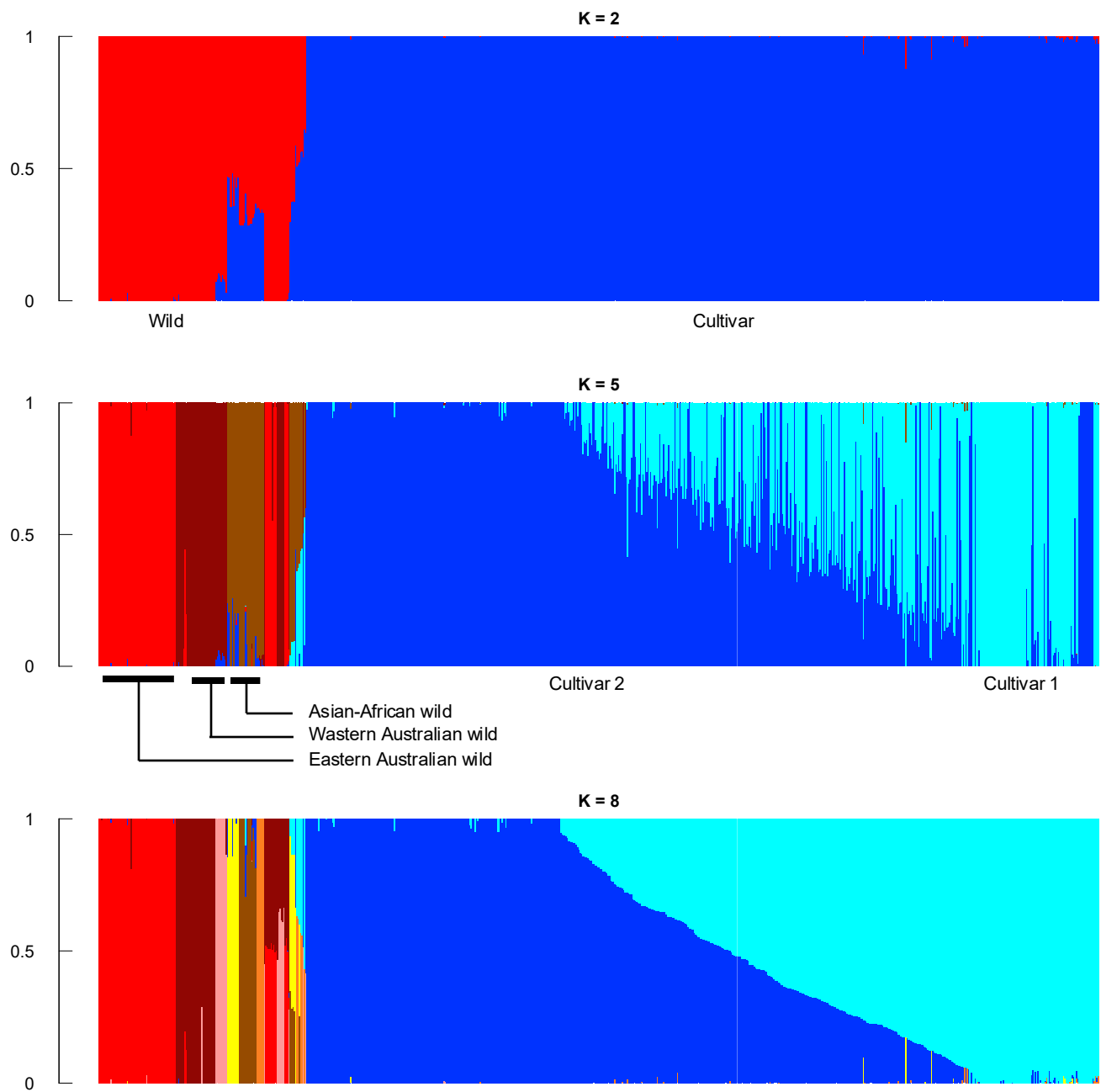

**Supplementary Fig. 6. Population structure of 780 mungbean accessions under K = 2, 5, and 8 in the ADMIXTURE analysis.** All individuals have the same order in the three panels. Population labels for K=8 are available in Figure 1a.

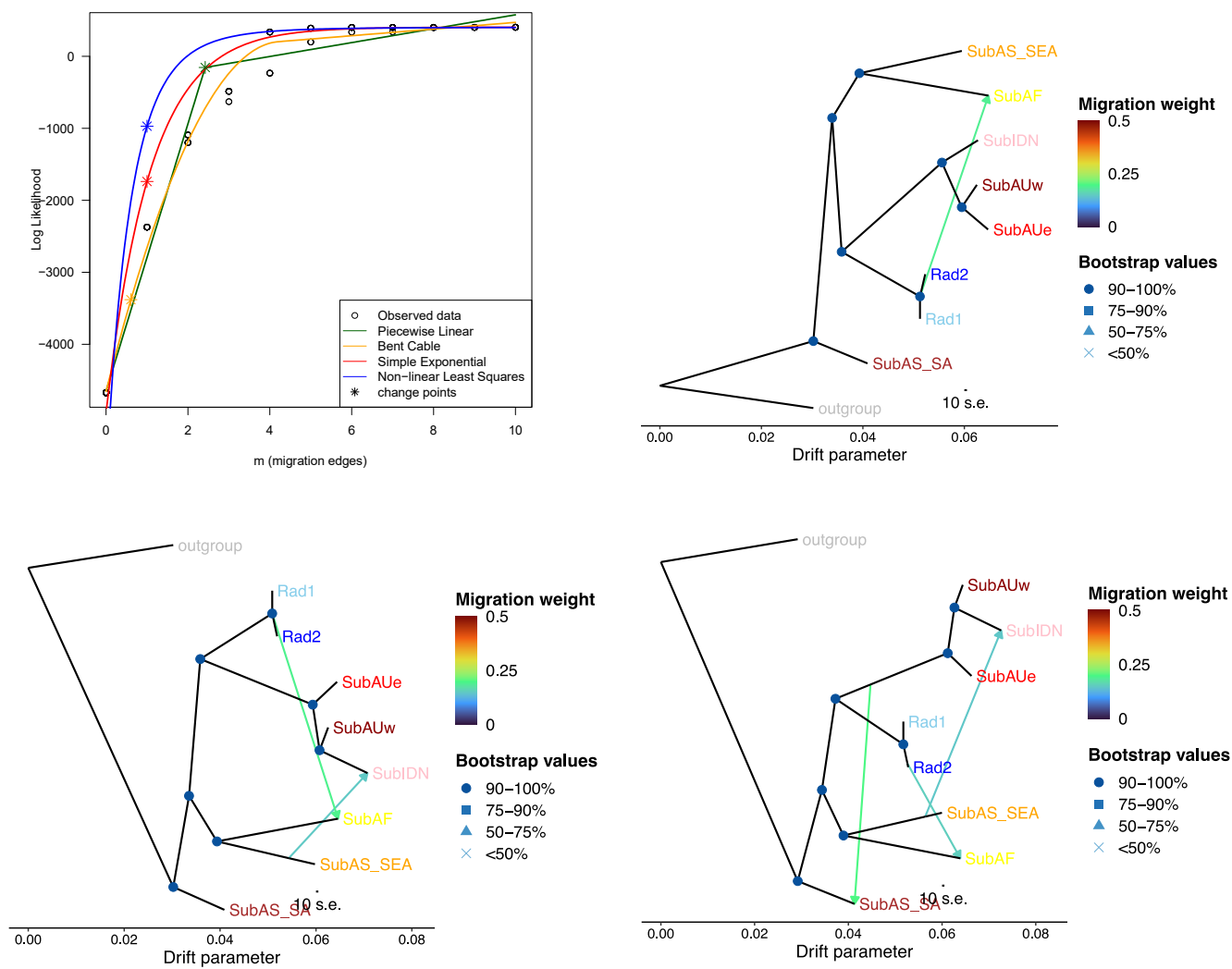

**Supplementary Fig. 7. Detailed Treemix results.** The OptM result on the upper left denotes the Log likelihood values of the potential number of migration events. The other plots are Treemix phylogeny assuming 1, 2, or 3 migration events.

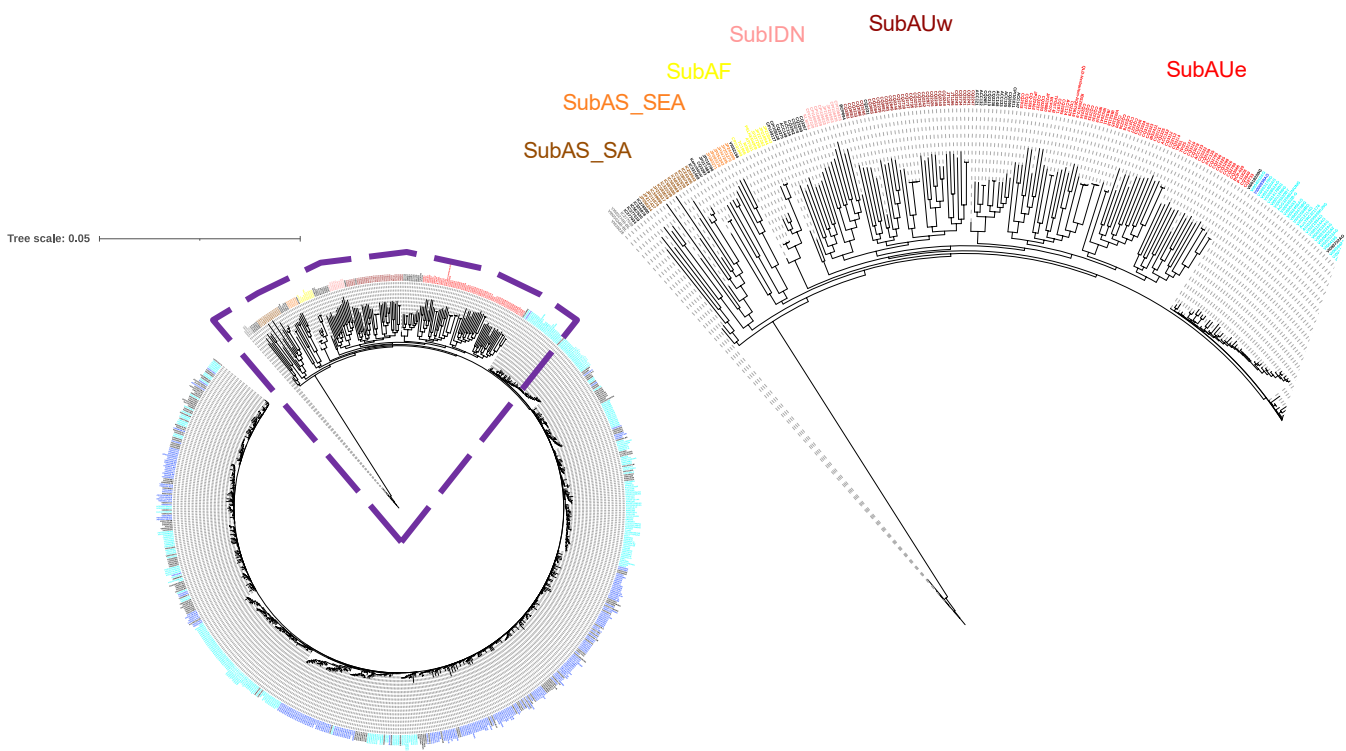

**Supplementary Fig. 8. Individual-based maximum likelihood tree of the nuclear genome of 780 mungbean accessions.** Four *V. mungo* accessions were used as the outgroup. The color of the text was based on ADMIXTURE K=8 in Fig. 1a. On the right is the zoom-in view of part of the overall tree, showing the relationship between wild and cultivar groups.

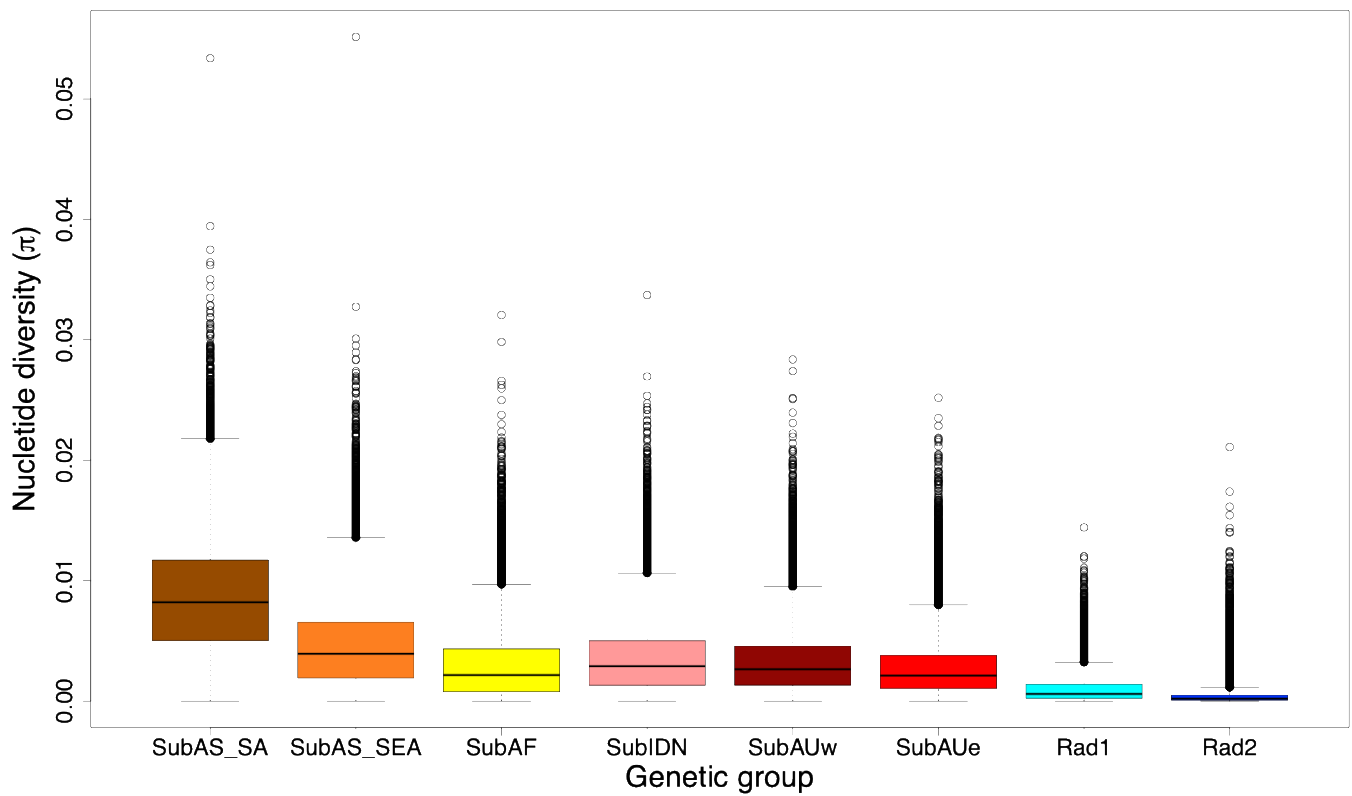

**Supplementary Fig. 9. Nucleotide Diversity of the eight genetic groups.** The boxes indicate medians and interquartile ranges, the whiskers indicate 95% values, and additional points in each boxplot represent outliers. Window size was set to 10kb.

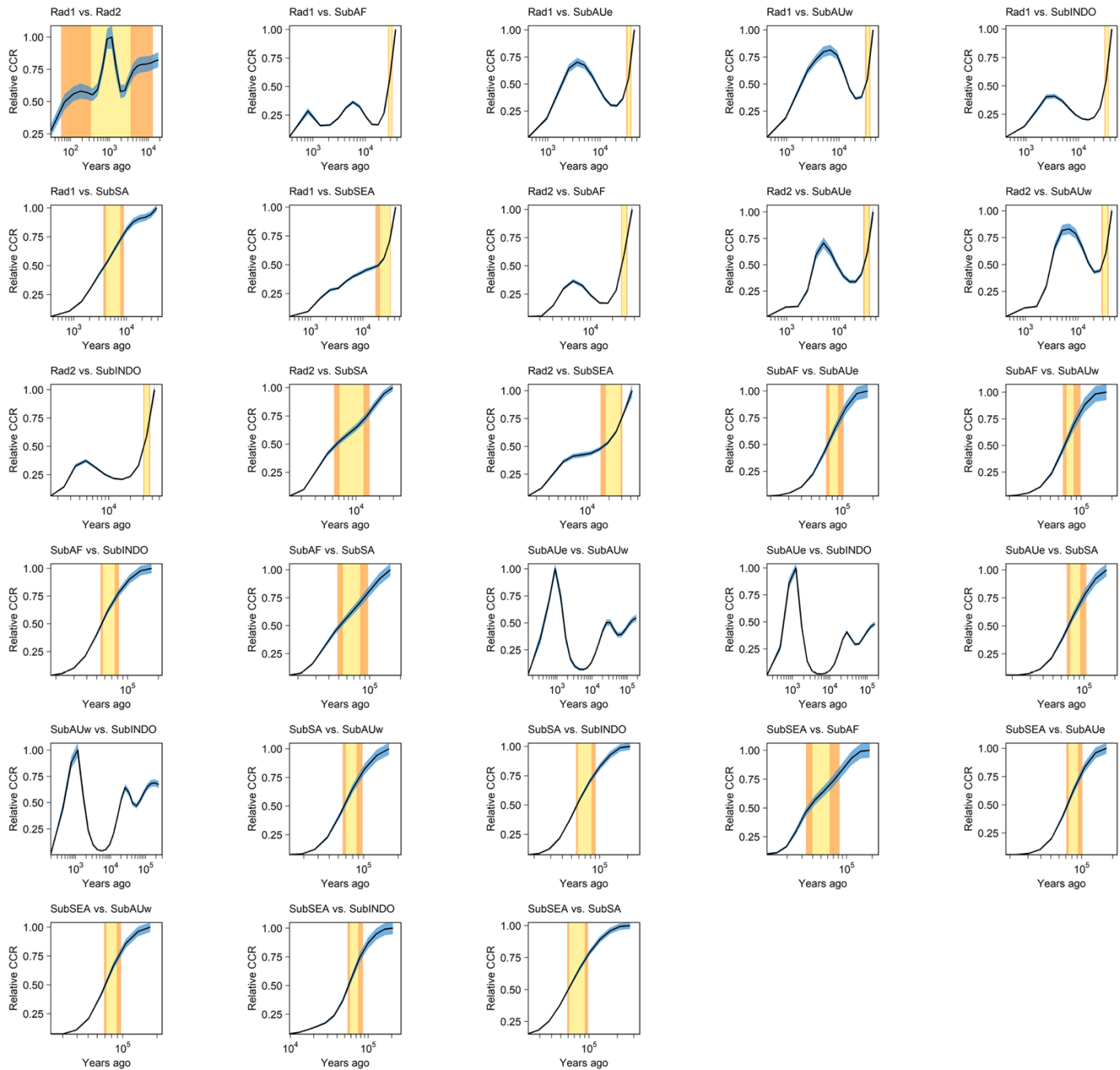

**Supplementary Fig. 10. Pairwise relative cross coalescence rate (RCCR) between population pairs estimated from MSMC.** One hundred repeats were conducted, and the range was shown as the blue shade. The orange bands denote the 95% range when RCCR reached 0.75 or 0.5, and the yellow bands denote the range between them.

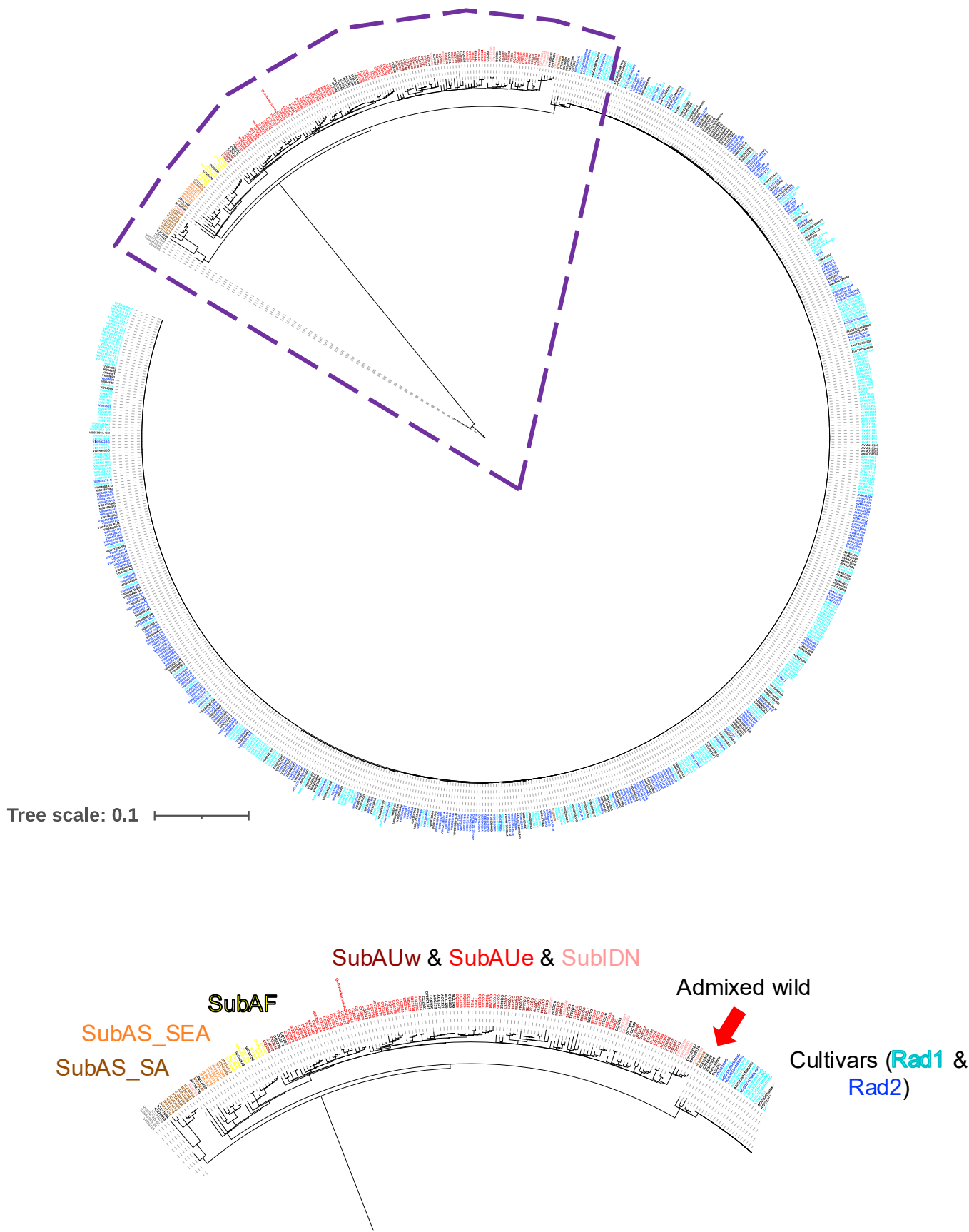

**Supplementary Fig. 11. Individual-based neighbor-joining tree of the chloroplast genome of 780 mungbean.** Four *V. mungo* accessions were used as the outgroup. The color of the text was based on ADMIXTURE K=8 in Fig. 1a. On the right is the zoom-in view of part of the overall tree, showing the relationship between wild and cultivar groups.

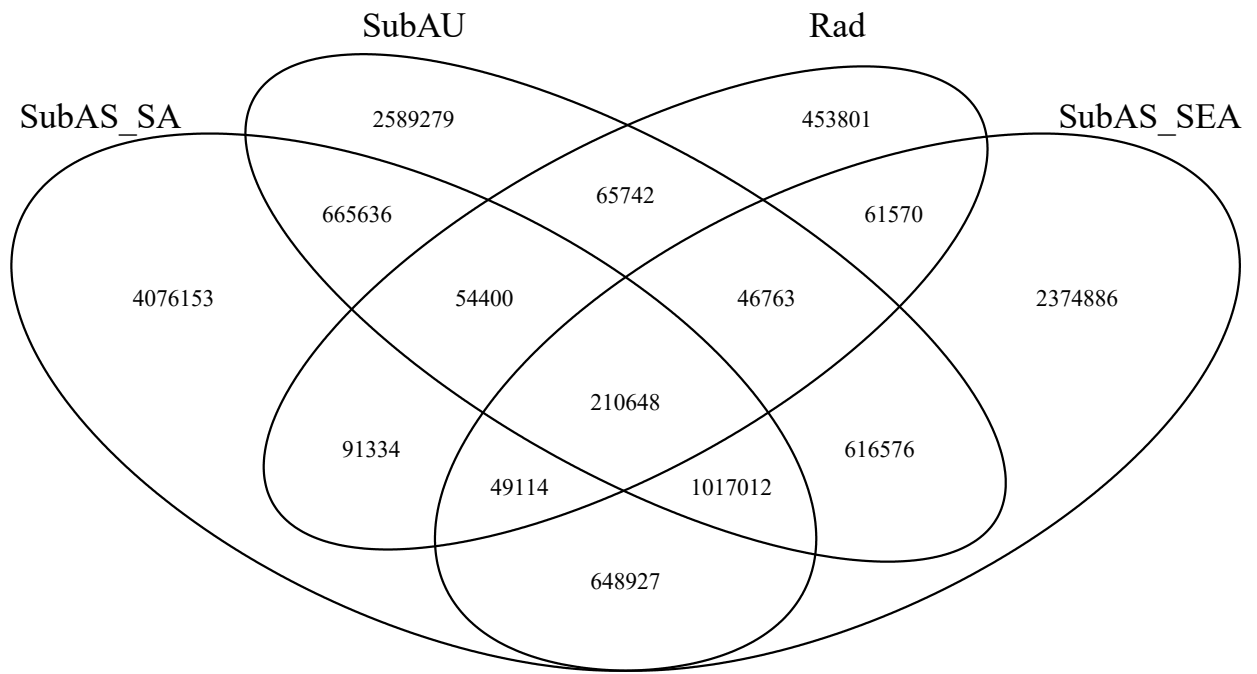

• SubAS\_SA: 6,813,224
• SubAU: 5,266,056
• SubAS\_SEA: 5,025,496
• Rad: 1,033,372
• SubAS\_SA ∩ Rad: 405,496
• SubAS\_SEA ∩ Rad: 368,095
• SubAU ∩ Rad: 377,553

**Supplementary Fig. 12. Number of SNPs private or shared among genetic groups.** The number of SNPs was calculated using a subset of 6 random accessions from each group to control for sample size differences.

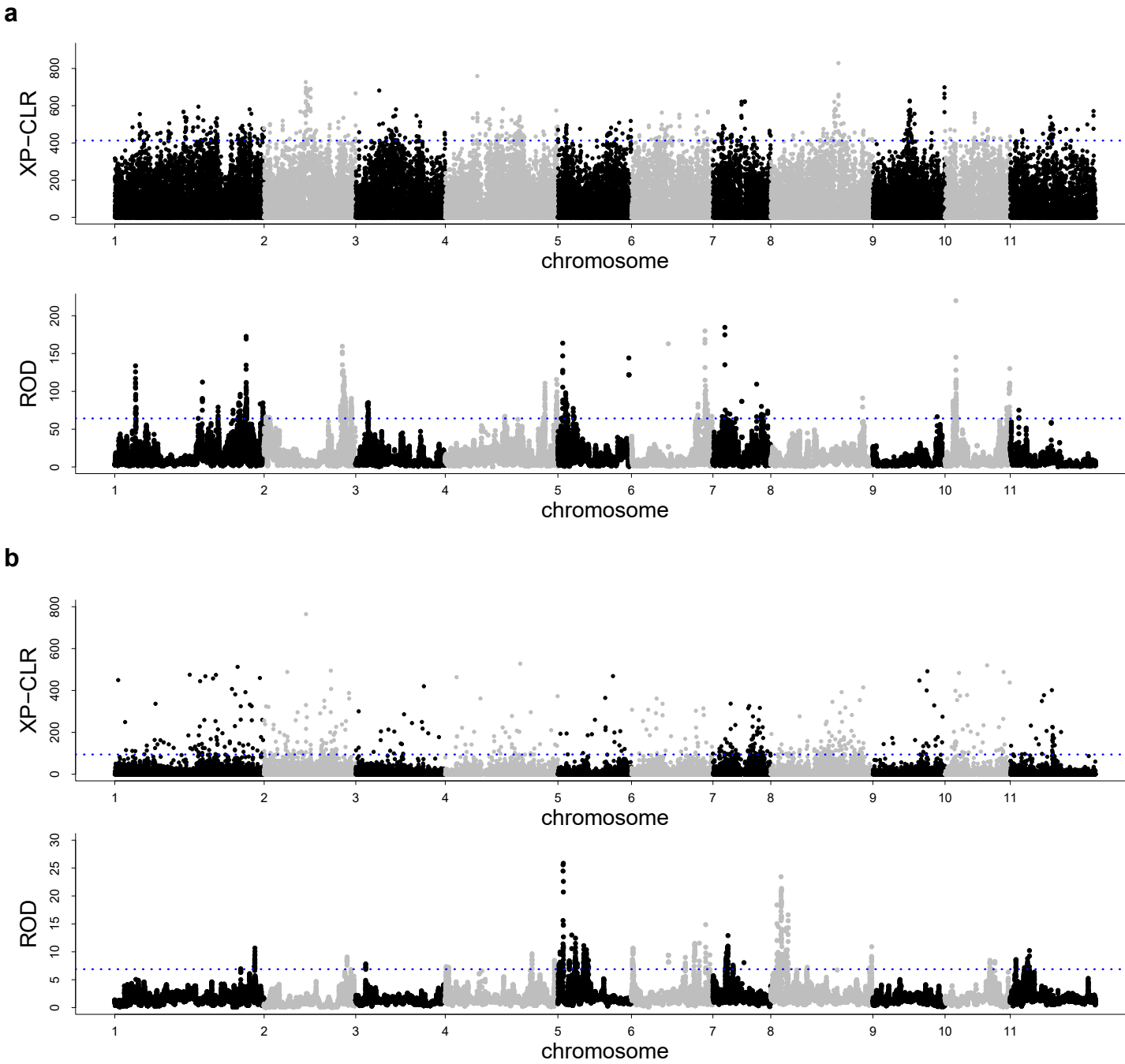

**Supplementary Fig. 13. Genome-wide selection scan of cross-population composite likelihood ratio (XP-CLR) and reduction of diversity (ROD) values.** Shown are the comparisons **a**, between wild and landrace and **b**, between landrace and improved cultivar lines. Window size was set to 100kb, and step size was set to 10kb. Blue dashed line: top 5 % threshold.

**a**

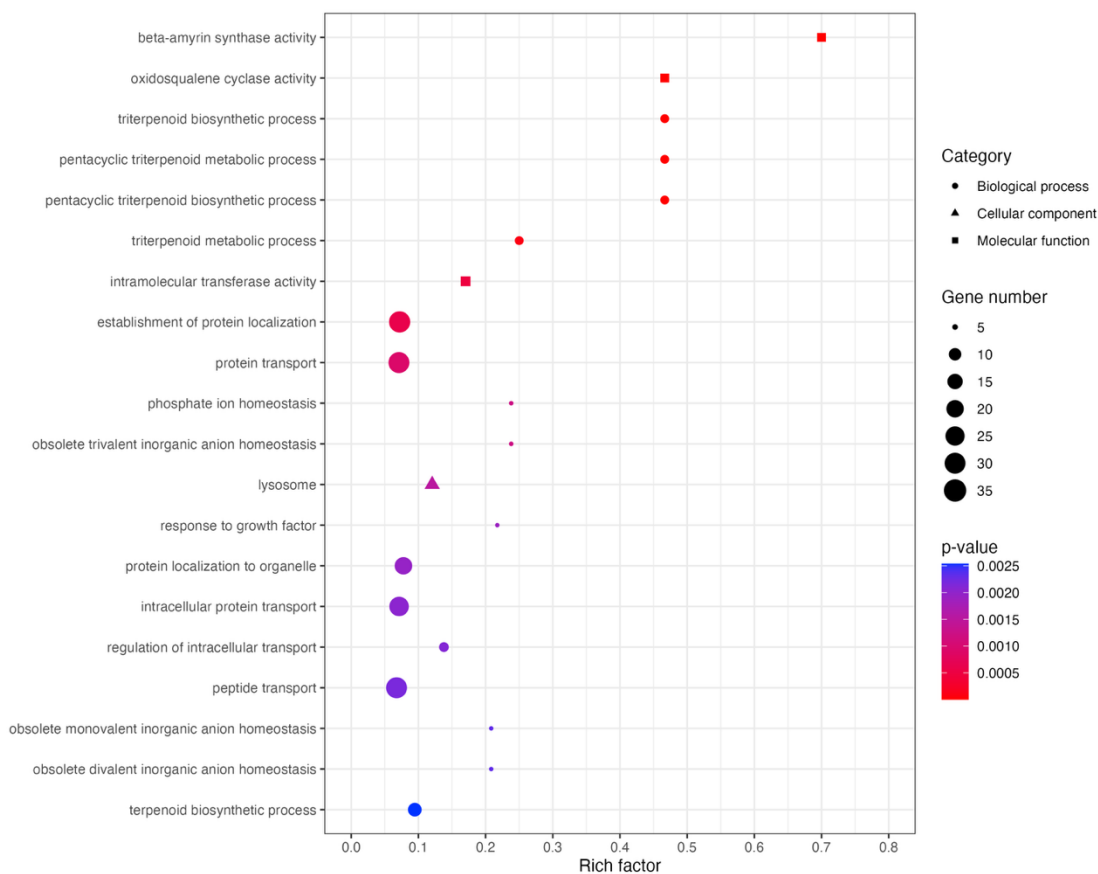

**b**

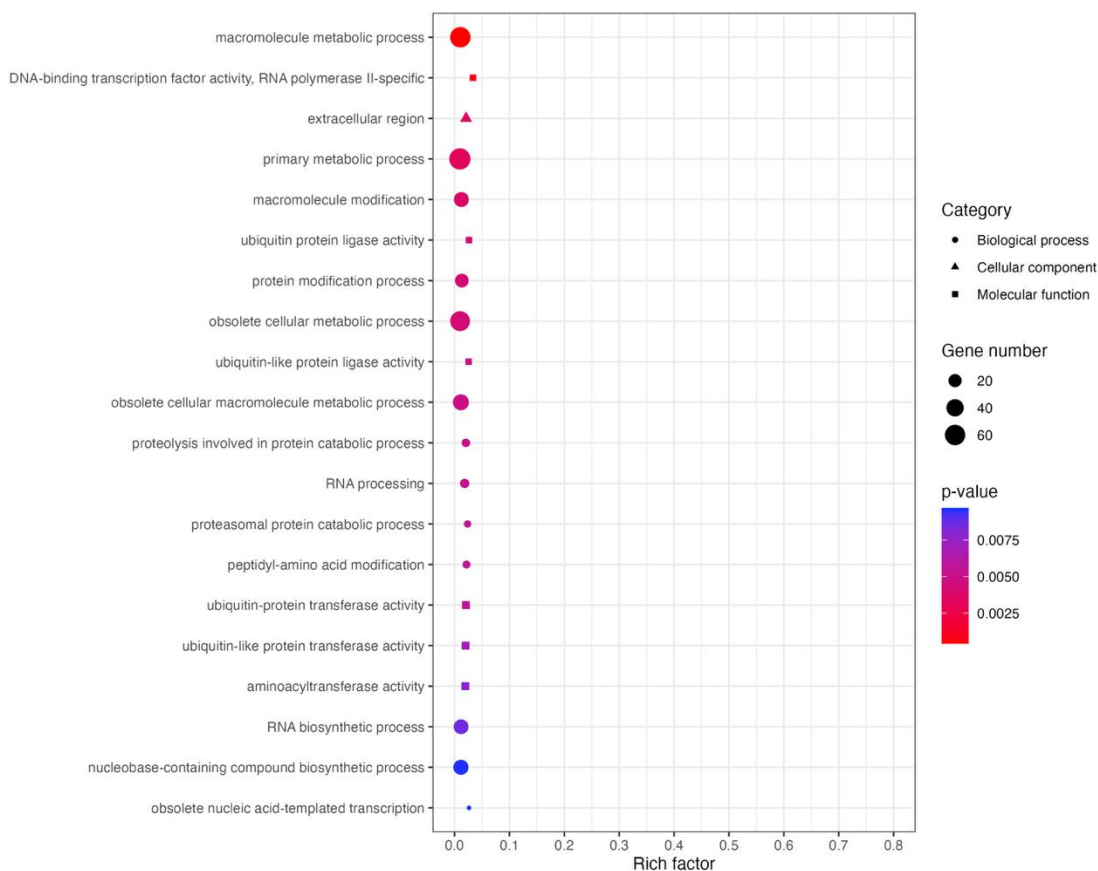

**Supplementary Fig. 14. Top 20 Gene Ontology (GO) terms enriched among candidate genes located in regions showing signatures of selection.** Shown are the GO enrichment results of candidate selective sweep genes from **a**, wild vs. landrace, and **b**, landrace vs. improved cultivar lines.

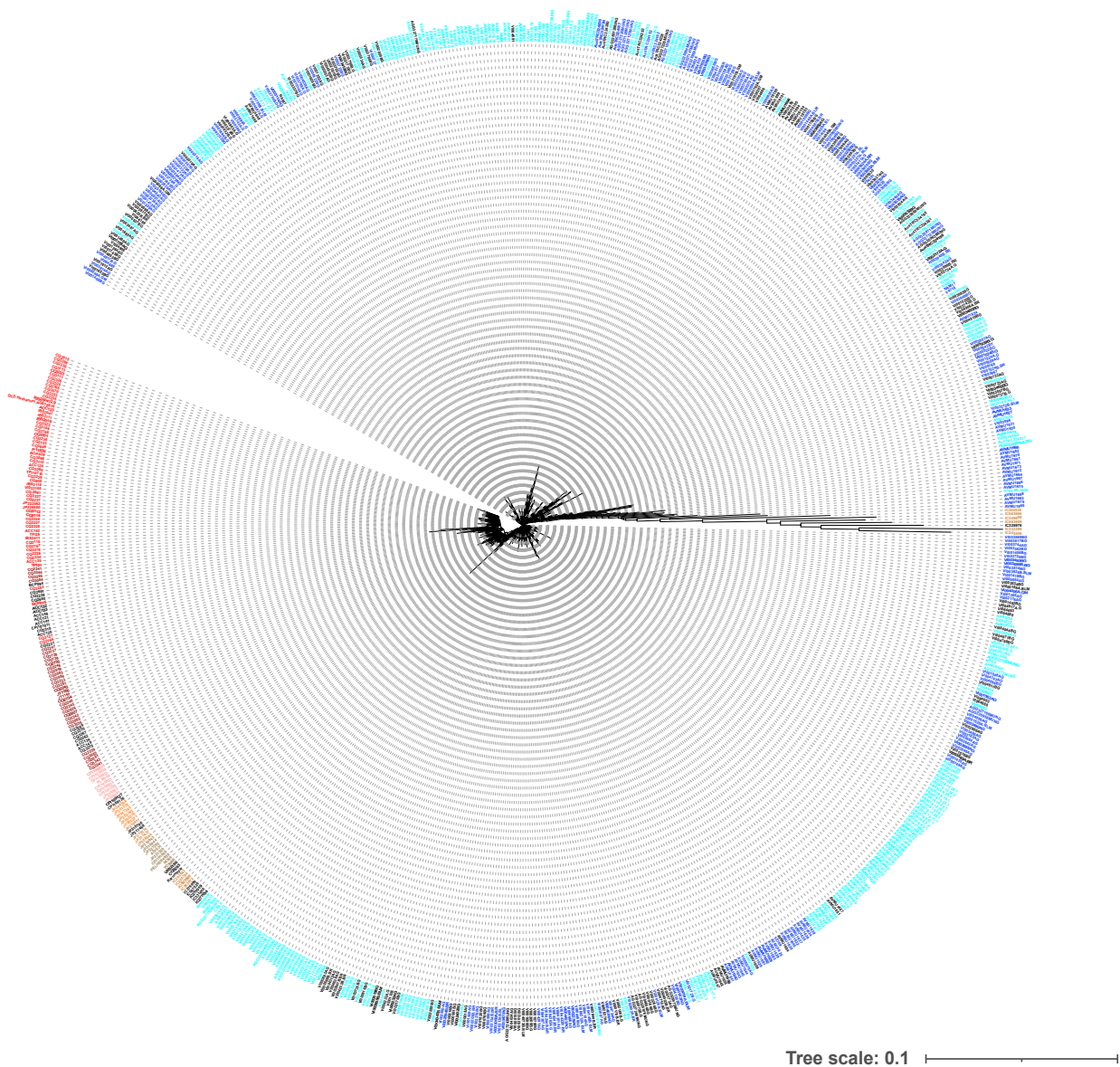

**Supplementary Fig. 15. Neighbor-joining tree of all mungbean accessions using the gene presence-absence variation dataset.**

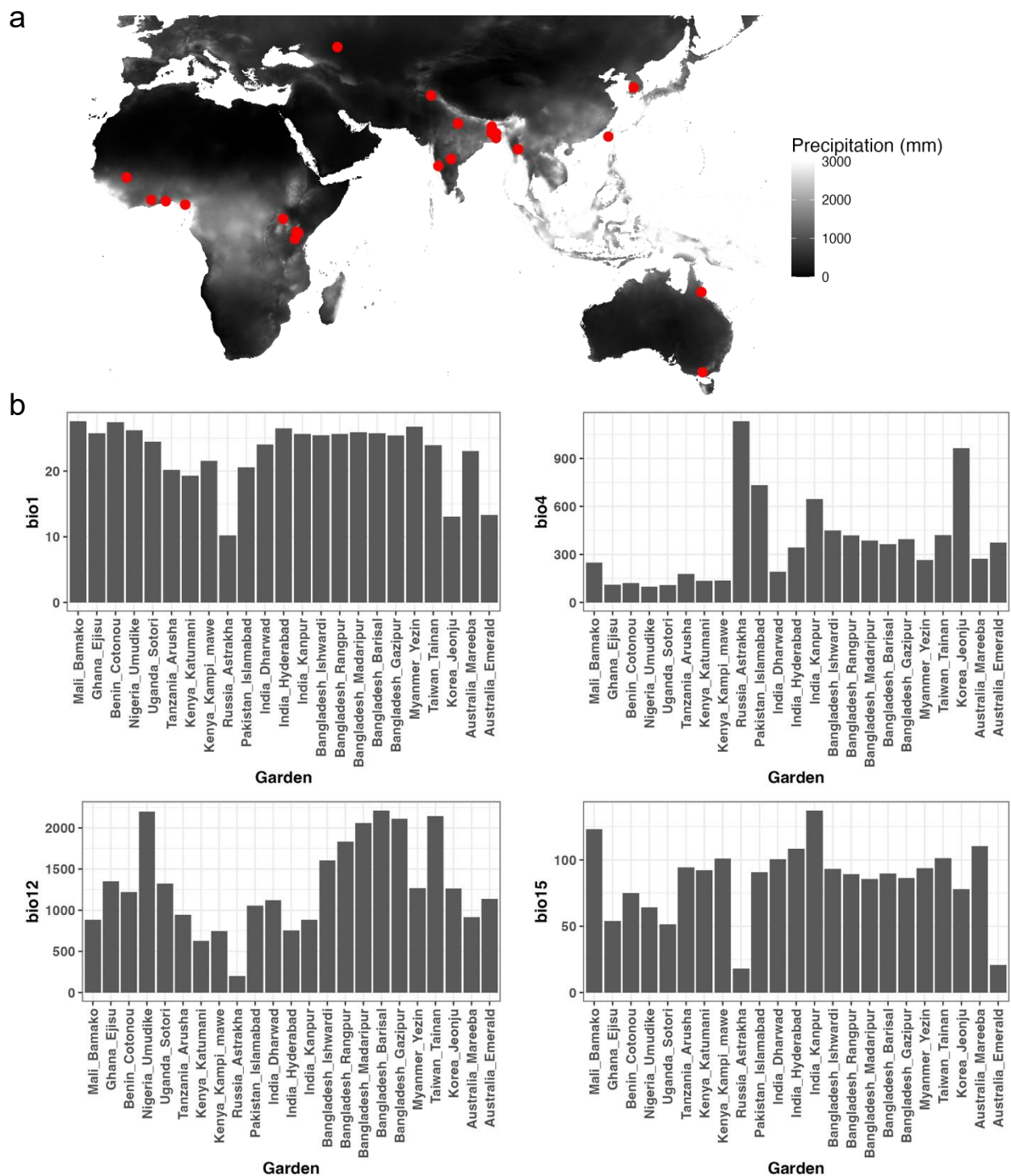

**Supplementary Fig. 16. Field trial sites of the International Mungbean Improvement Network (IMIN). a**, Twenty-three field trial sites. **b**, Variation in annual mean temperature (bio1), temperature seasonality (bio4), annual precipitation (bio12), and precipitation seasonality (bio15).

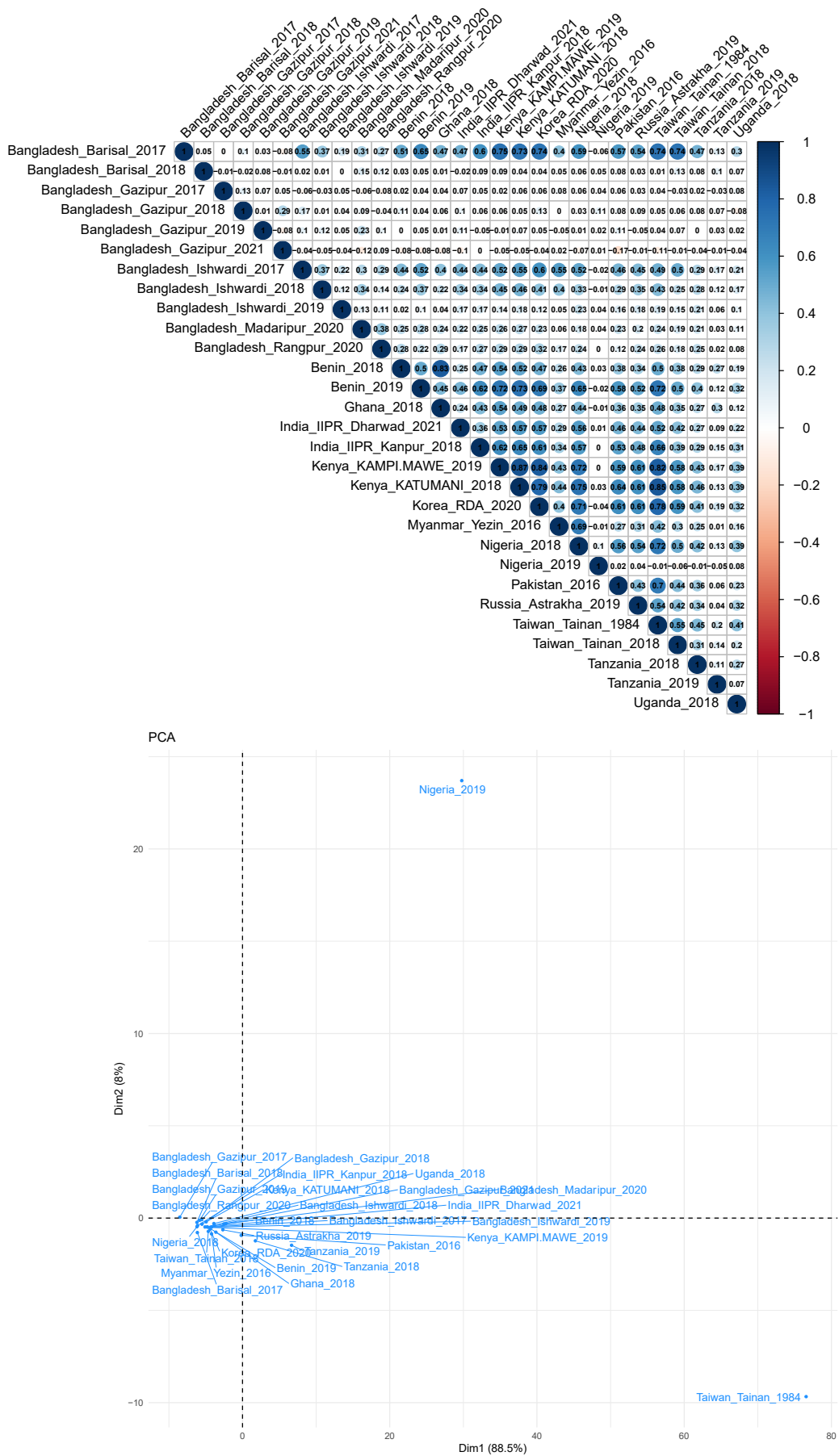

**Supplementary Fig. 17. Pairwise correlation of seed weight across trials and the among-trial principal component analysis.**

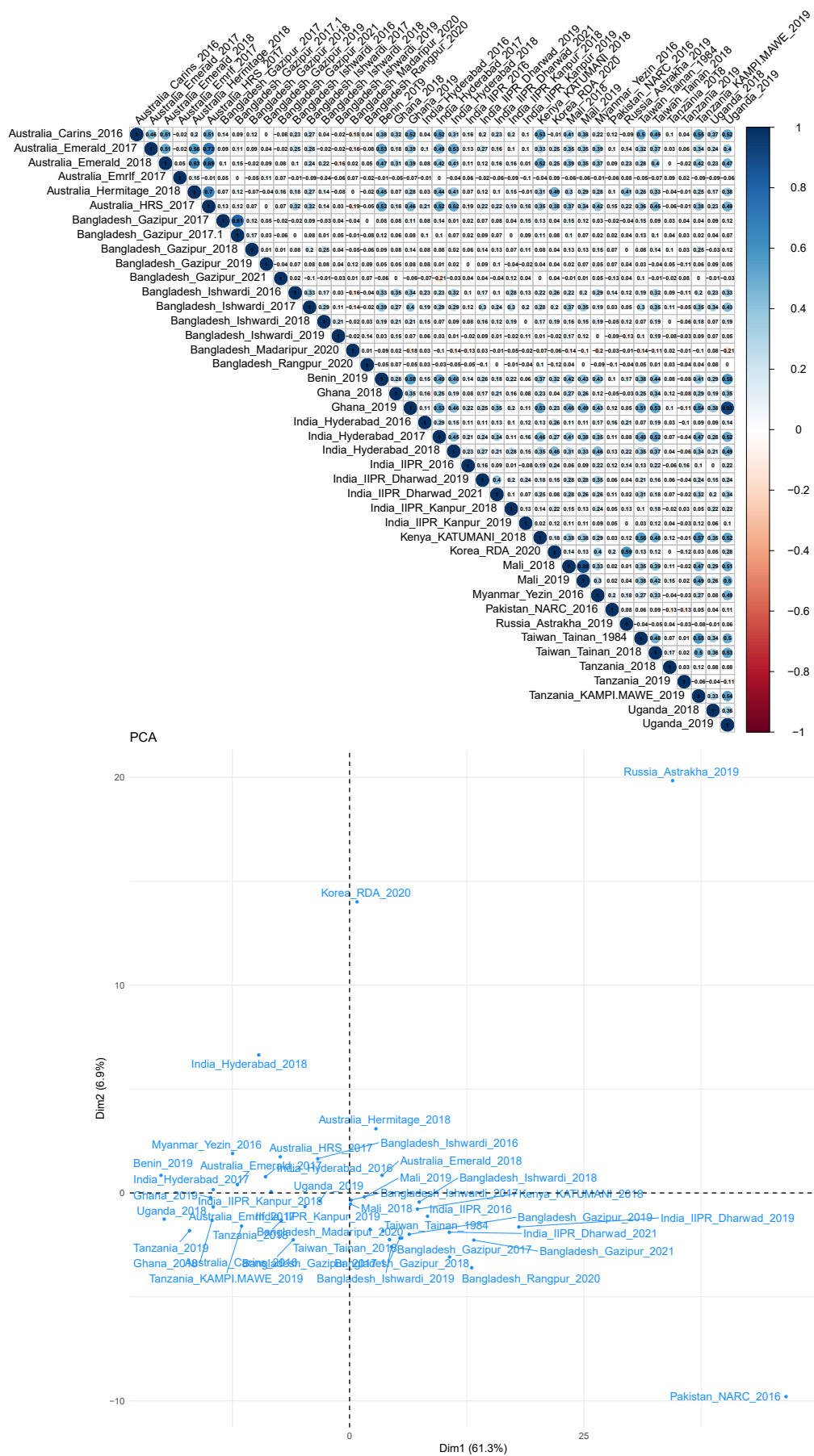

Supplementary Fig. 18. Pairwise correlation of days to flowering across trials and the among-trial principal component analysis.

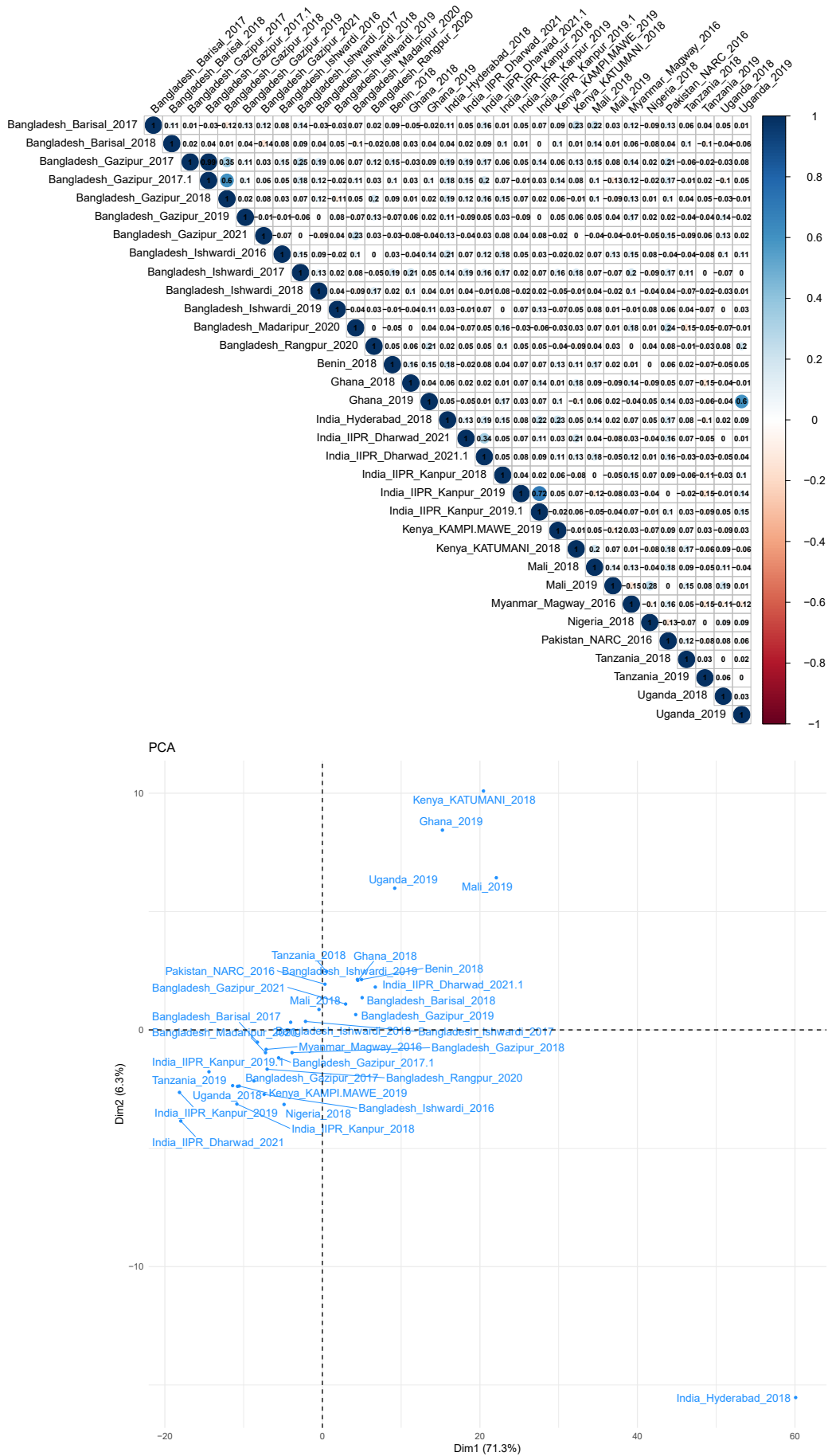

**Supplementary Fig. 19. Pairwise correlation of number of pods per plant across trials and the among-trial principal component analysis.**

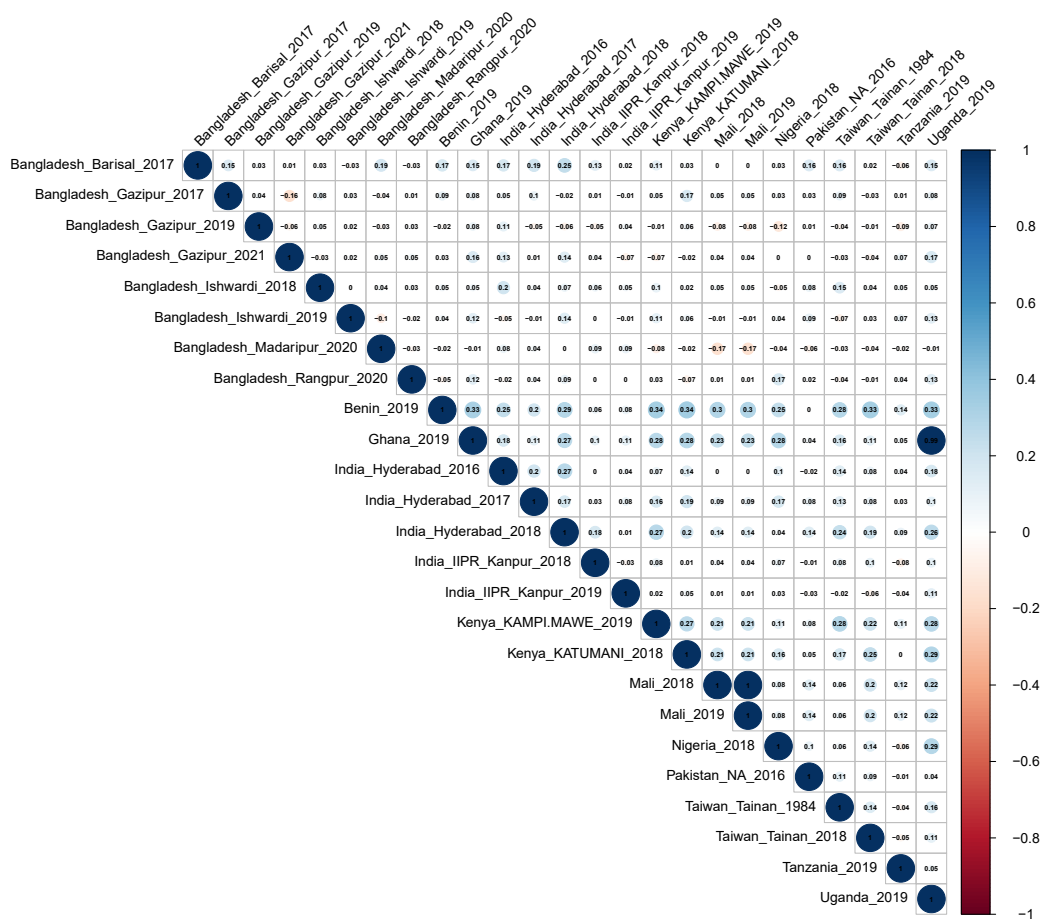

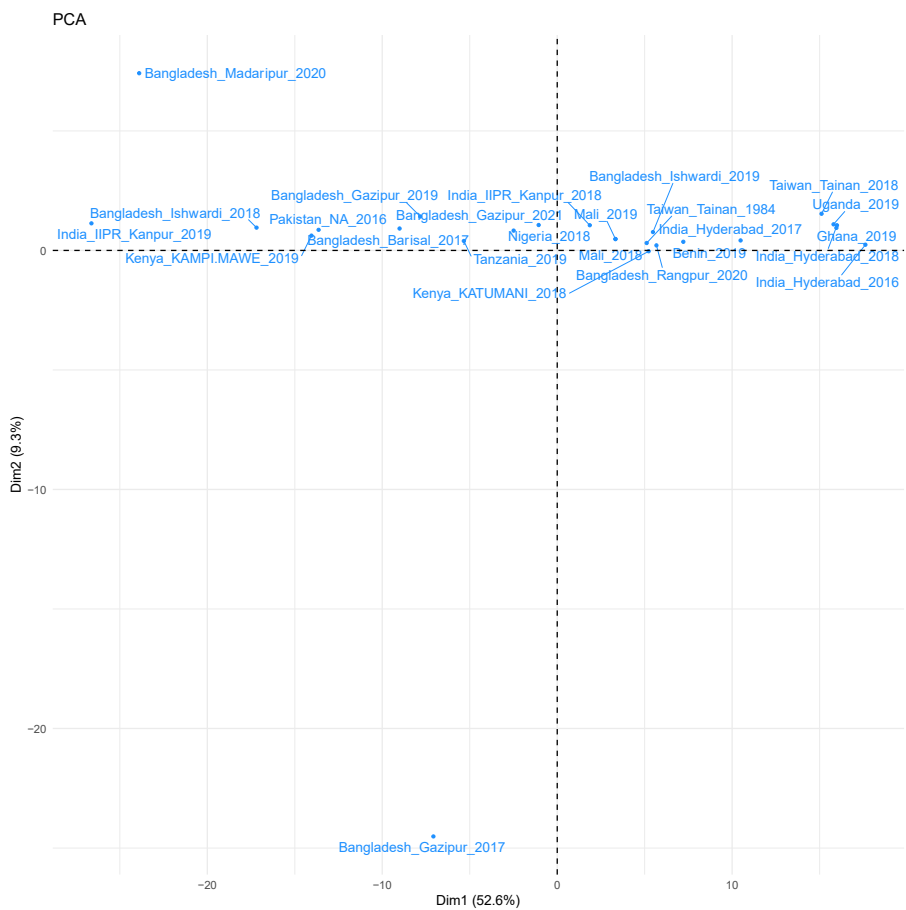

**Supplementary Fig. 20. Pairwise correlation of number of seeds per pod across trials and the among-trial principal component analysis.**

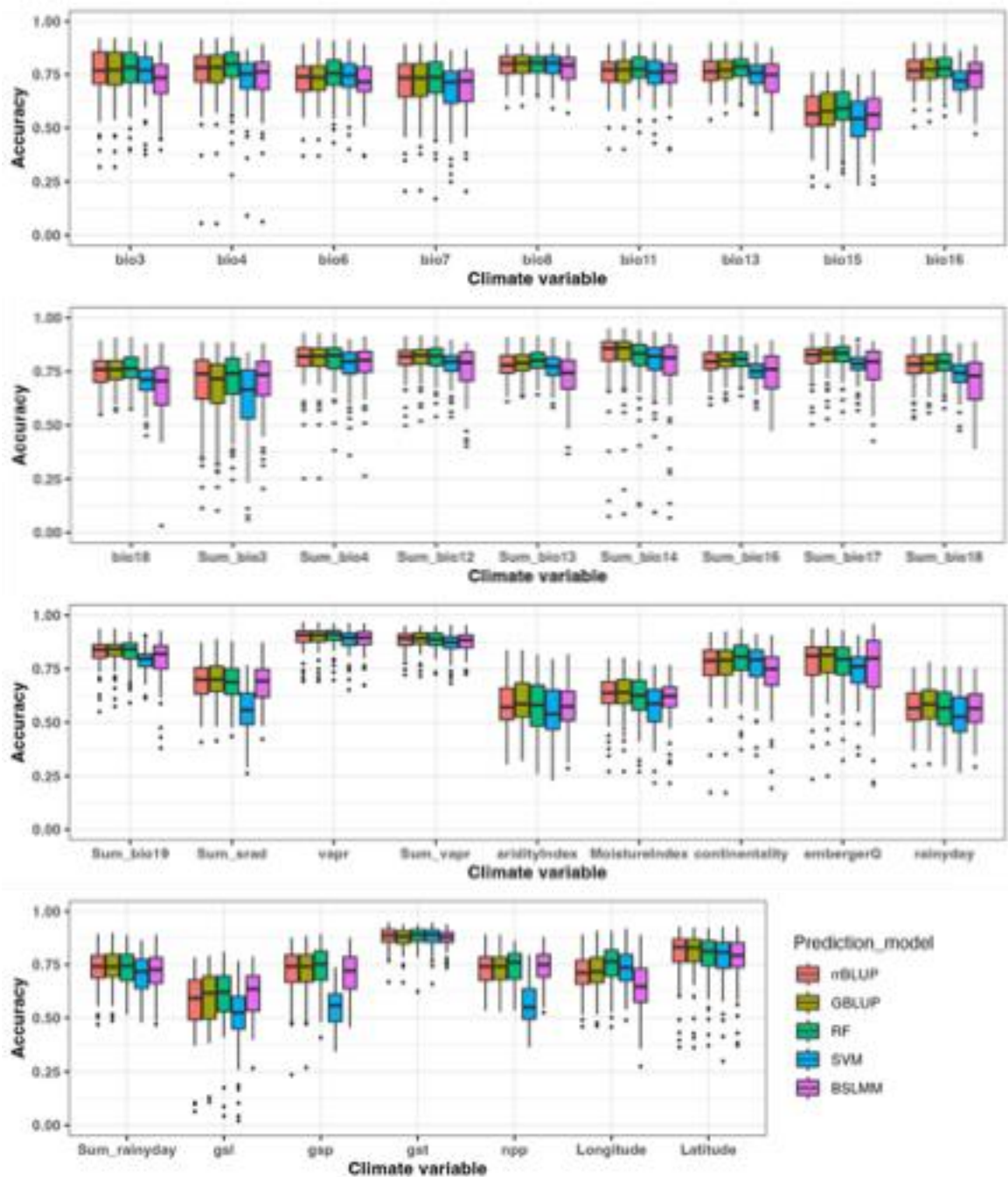

**Supplementary Fig. 21. Accuracy of genomic prediction of 34 climate variables using 5 different genomic prediction models.**

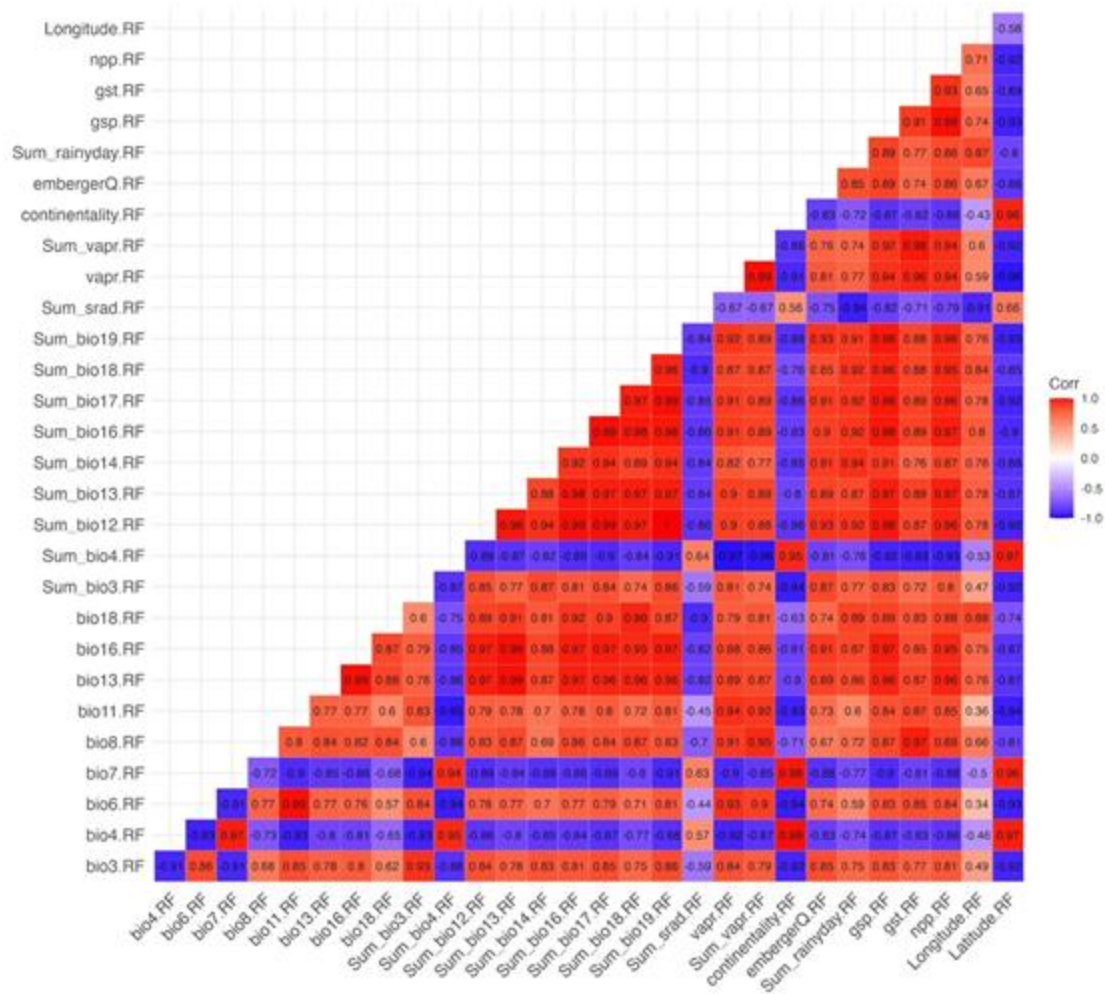

Supplementary Fig. 22. Correlation heatmap of 29 predicted climate variables.

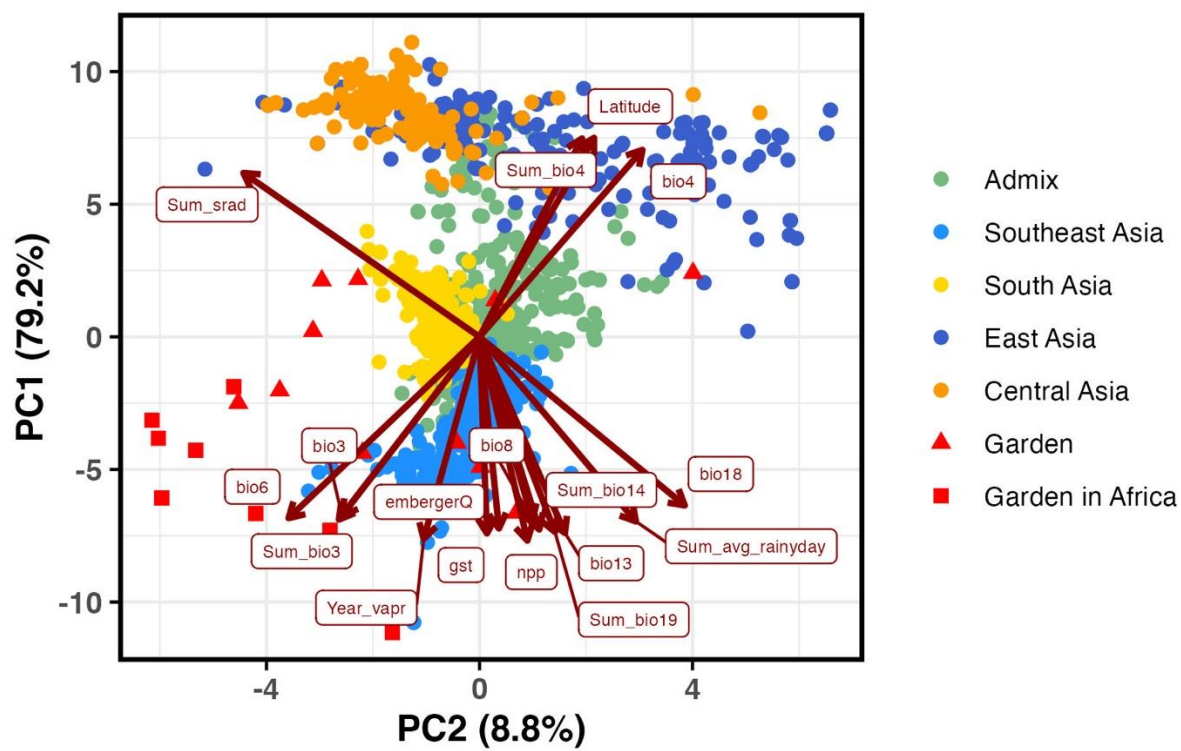

**Supplementary Fig. 23. Principal component analysis (PCA) biplot of 17 predicted native climate variables ($r < 0.95$).** Arrows represent the direction and strength of each variable's contribution to the first two principal components (PC1 and PC2).

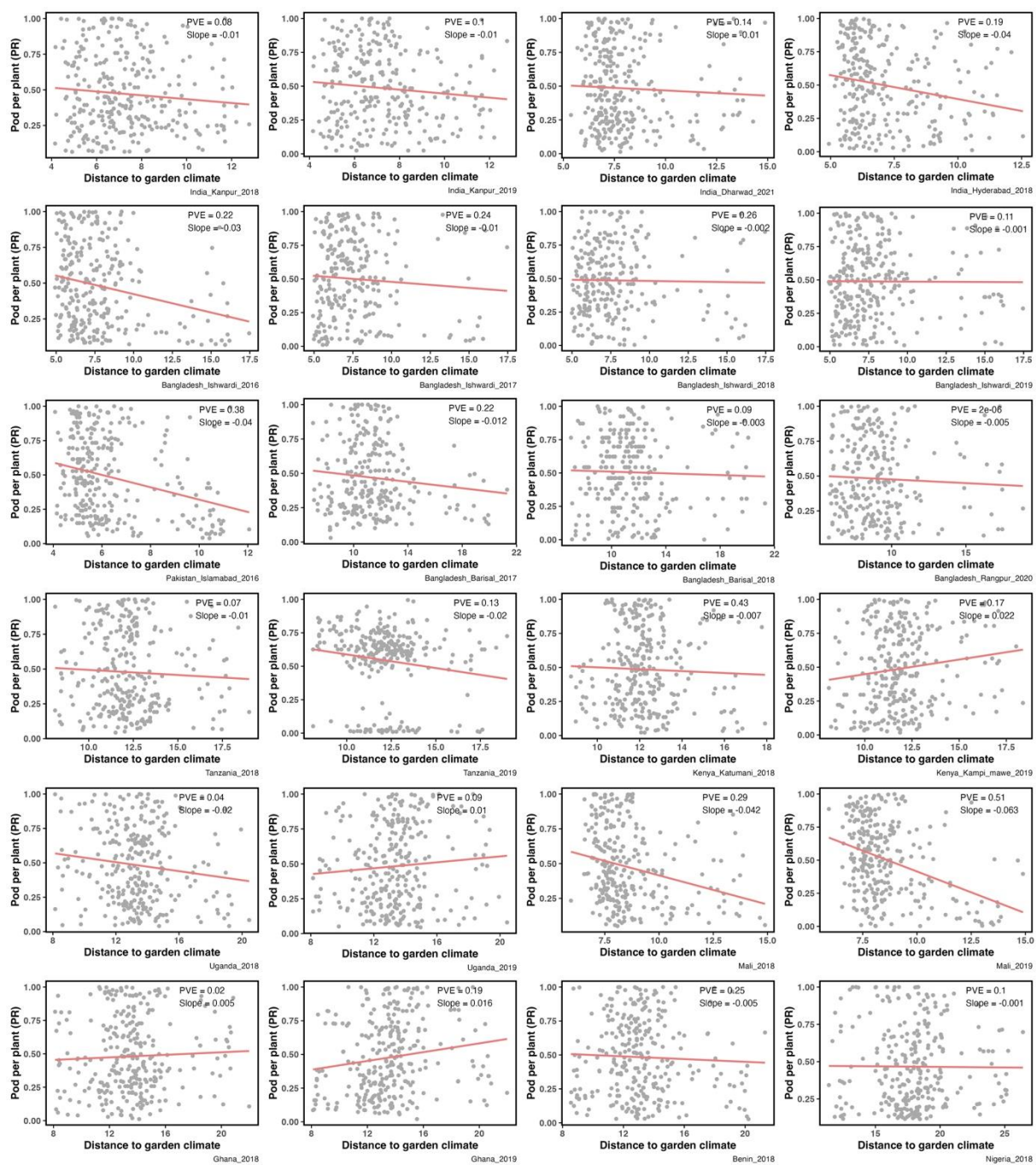

**Supplementary Fig. 24. Scatter plots showing the relationship between the percentile rank (PR) of number of pod per plant and “the climate distance between each accession’s native environment and the garden environment”. PVE denotes genome-estimated heritability.**

a

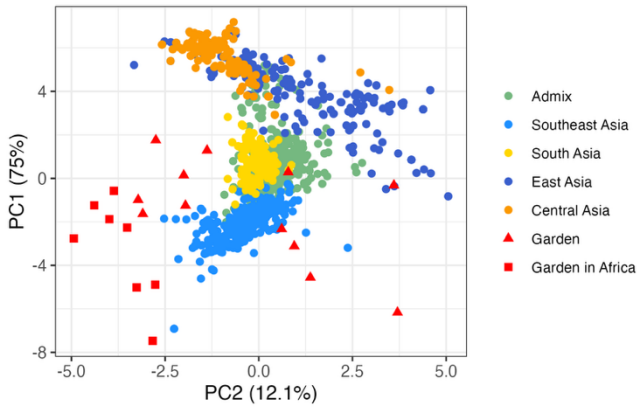

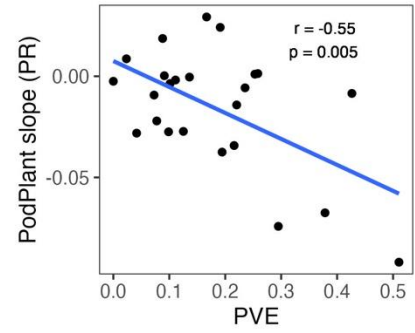

b

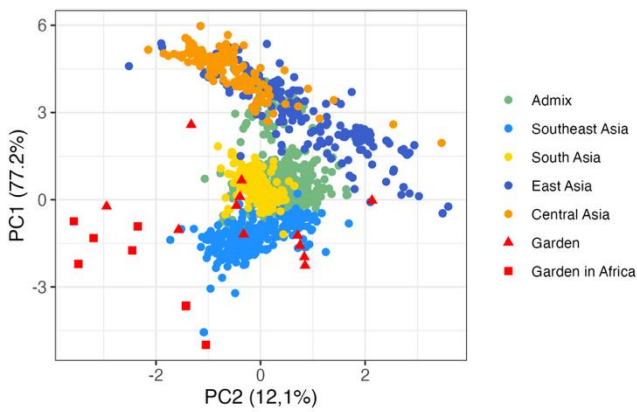

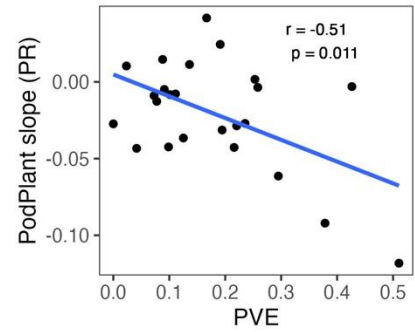

**Supplementary Fig. 25. Comparison of climate variable sets selected under different correlation thresholds.** Climate variable sets were filtered at **a**, $r < 0.9$ and **b**, $r < 0.8$, resulting in 7 and 4 variables, respectively. Despite the reduction in the number of variables, PCA results show similar distribution patterns, and the relationship between trait heritability (PVE) and yield response slopes remains consistent across thresholds.

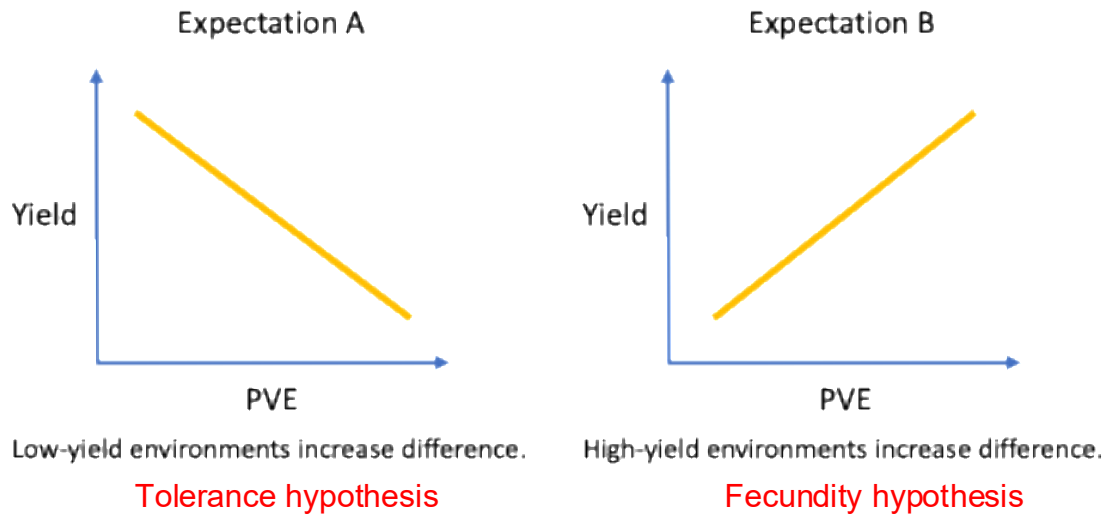

**Supplementary Fig. 26. Two hypotheses explaining why trial sites with higher heritability (PVE) also exhibit stronger patterns of accession local adaptation.** In the tolerance hypothesis, suboptimal growing conditions (lower yield) exert local adaptation and higher PVE by allowing tolerant accessions from similar environments to thrive, while most accessions performed well in benign conditions. In the fecundity hypothesis, benign conditions (higher yield) enable the potential of high-yield accessions, while most accessions performed less well in suboptimal conditions.
